# Environmental vulnerability of the global ocean plankton community interactome

**DOI:** 10.1101/2020.11.09.375295

**Authors:** S. Chaffron, E. Delage, M. Budinich, D. Vintache, N. Henry, C. Nef, M. Ardyna, A.A. Zayed, P.C. Junger, P.E. Galand, C. Lovejoy, A. Murray, H. Sarmento, Tara Oceans coordinators, S. Acinas, M. Babin, D. Iudicone, O. Jaillon, E. Karsenti, P. Wincker, L. Karp-Boss, M.B. Sullivan, C. Bowler, C. de Vargas, D. Eveillard

**Affiliations:** Université de Nantes, CNRS UMR 6004, LS2N, F-44000 Nantes, France; Research Federation (FR2022) Tara Océan GO-SEE, Paris, France; Sorbonne Université, CNRS, Laboratoire Adaptation et Diversité en Milieu Marin, Station Biologique de Roscoff, 29680 Roscoff, France; Institut de Biologie de l’École Normale Supérieure (IBENS), École Normale Supérieure, CNRS, INSERM, PSL Université Paris, 75005 Paris, France; Stanford University, Department of Earth System Science, Stanford, CA 94305, United States; Sorbonne Université, CNRS, Laboratoire d’Océanographie de Villefranche, LOV, F-06230, Villefranche-sur-Mer, France; Department of Microbiology, The Ohio State University, Columbus, OH 43210, United States; Department of Hydrobiology, Universidade Federal de São Carlos (UFSCar), Rodovia Washington Luiz, 13565-905 São Carlos, SP, Brazil; Sorbonne Université, CNRS, Laboratoire d’Ecogéochimie des Environnements Benthiques, LECOB, 66500, Banyuls-sur-Mer, France; Département de biologie, Faculté des sciences et Institut de biologie intégrative et des systèmes (IBIS) 1030, ave de la Médecine, Université Laval, Québec QC, Canada; Division of Earth and Ecosystem Science, Desert Research Institute, Reno, NV 89512, USA; Department of Marine Biology and Oceanography, Institut de Ciències del Mar (CSIC), Barcelona, 08003, Spain; Takuvik International Research Laboratory, Université Laval and CNRS, Québec QC, Canada; Stazione Zoologica Anton Dohrn, Villa Comunale, Naples, 80121, Italy; Génomique Métabolique, Genoscope, Institut François Jacob, CEA, CNRS, Université Evry, Université Paris-Saclay, Evry, France; School of Marine Sciences, University of Maine, Orono, ME, USA; Department of Civil, Environmental and Geodetic Engineering, The Ohio State University, Columbus, OH 43210, United States

## Abstract

Marine plankton form complex communities of interacting organisms at the base of the food web, which sustain oceanic biogeochemical cycles, and help regulate climate. Though global surveys are starting to reveal ecological drivers underlying planktonic community structure, and predicted climate change responses, it is unclear how community-scale species interactions will be affected by climate change. Here we leveraged *Tara* Oceans sampling to infer a global ocean cross-domain plankton co-occurrence network – *the community interactome* – and used niche modeling to assess its vulnerabilities to environmental change. Globally, this revealed a plankton interactome self-organized latitudinally into marine biomes (Trades, Westerlies, Polar), and more connected poleward. Integrated niche modeling revealed biome-specific community interactome responses to environmental change, and forecasted most affected lineages for each community. These results provide baseline approaches to assess community structure and organismal interactions under climate scenarios, while identifying plausible plankton bioindicators for ocean monitoring of climate change.

## Introduction

Marine plankton and associated processes are at the core of global biogeochemical cycles, shaping ecosystem structure and influencing climate regulation (*1*). While global biodiversity maps for viruses, prokaryotes and microbial eukaryotes are beginning to emerge (*2-4*), identifying and understanding the complex network of interactions between these organisms and their environment is in its infancy (*5*). These interactions are critical to establish the ecosystem trophic links that underpin biogeochemical cycles and feedbacks that drive climate regulation and response (*6, 7*). While abiotic factors, such as temperature, can explain a large fraction of microbial community composition in the global ocean (*8*), biotic interactions can differentially shape ecosystem diversity (*9*), and can even influence the adaptation to new environments (*10*). Thus, determining how plankton ecological interactions are structured and affected by environmental change remains a significant challenge.

Large-scale holistic marine ecosystem sampling facilitates conceptualization of plankton community interactomes as co-occurrence networks that are useful to model the complex community structure of ecological associations (*11, 12*). Such networks have enabled the detection of communities assembled through niche overlap across biomes (*13*), and also the prediction of putative interactions such as parasitism or symbioses (*14*). Likewise, plankton co-occurrence networks have been instrumental in detecting interrelated changes in community structure from surface to depth (*15*), as well as to identify specific communities of key lineages (e.g. *Synechococcus*, its phages, and Collodaria) associated with global open ocean processes such as carbon export (*16*). Community interactomes are also useful to identify central, highly connected lineages that may play significant ecological roles and confer stability to the community (*17*). These central lineages can correspond to keystone taxa that are good indicators of community shifts (*18*). Understanding the mechanisms affecting these central taxa may help us to predict responses of microbiome structure and functioning to perturbations (*19*).

While community interactomes inferred from global-scale samplings summarize well the complexity and potential interactions within microbial assemblages (*12*), they usually do not reflect dynamic processes shaping the observed system, as measured by longitudinal high-frequency sampling (*20*). Thus, alternative strategies need to be developed to capture ecosystem dynamics and responses from spatial samplings. Indeed, plankton species display various ecological and evolutionary responses to global environmental change (*21, 22*). Within marine ecosystems, the interplay between species ecological niche and climate change can induce abrupt community shifts, which may lead to long-term reconfiguration of marine metazoan communities or biodiversity rearrangements (*23, 24*). Recently, environmental drivers of ocean plankton diversity were inferred from *Tara* Oceans data and used to predict the effects of severe warming on surface ocean biodiversity (*3*). While species niche distribution models combined with climate models are useful to project fine-scale future distributions of species (*25, 26*), species interactions are generally not included in these models (*27*), certainly due to our lack of knowledge about organismal interactions. Nevertheless, plankton network metrics can capture emergent properties (e.g. connectivity) relating to ecological characteristics of the community (*28*), which can serve as proxies of ecosystem and community-level resilience (*29*). Given that biotic interactions can influence species distributions at macroecological scales (*30*), and that climate change may cause trophic cascading effects on plankton community structure by directly impacting the top and bottom of marine food webs (*31*), ecological interactions need to be considered for assessing plankton community stability under climate change scenarios.

Here, we bridge the gap between ecological and climate modeling by combining network analyses (*32*) with species niche models (*33*) into a novel computational framework for predicting ecosystem-scale vulnerabilities to environmental change. By leveraging *Tara* Oceans data from all major oceanic provinces, including the Arctic Ocean, we inferred a comprehensive global-ocean cross-domain plankton co-occurrence network from sequencing data. We built statistical niche models to predict realized niche widths of planktonic taxa, across kingdoms, and from pole-to-pole. These were then mapped onto the network and used to evaluate both local and global robustness of plankton community structures to simulated environmental changes. Noticeable efforts have used the ecological niche concept to identify open ocean physical conditions governing phytoplankton biogeography (*34*), and also to better formalize central biogeochemical processes through the definition of key plankton functional types (*35*). The niche representation of planktonic diversity affords a more effective integration of abiotic and biotic constraints to better predict perturbations of primary productivity under climate change scenarios (*26, 36*).

## Results and Discussion

### A cross-kingdom plankton interactome from pole to pole

To reconstruct a global marine plankton co-occurrence network across kingdoms of life, we analyzed data from 115 stations from the *Tara* Oceans expeditions (2009-2013) covering several organismal size fractions and all major oceanic provinces (*37*) across an extensive latitudinal temperature gradient from pole to pole (Fig. 1A). Via a dedicated probabilistic learning algorithm (*38*) (see M&M), we predicted ecological interactions between plankton taxa from compositional abundances inferred from sequencing data. The resulting integrated species association network (referred to as the *Global Ocean Plankton Interactome –* GPI) counts a total of 20,810 nodes corresponding to Operational Taxonomic Units (OTUs) and 86,026 edges corresponding to potential biotic interactions (Fig. 1A). In comparison to a previous plankton interactome generated from *Tara* Oceans data (*14*), GPI doubled the number of recovered known interactions from the literature (see supplementary text). A vast majority of positive associations (98.5%) were predicted, probably underlying a prevalent role for biotic interactions in shaping marine plankton communities (*14*). Very few direct associations between OTUs and environmental parameters were detected (n = 325, see supplementary text). However, by estimating robust ecological optima (or niche value) and tolerance ranges (or niche widths) (*39*) for each OTU and environmental parameter (see M&M and Table S3), we observed a strong influence of temperature in structuring predicted interactions (Fig. 1A). The GPI displayed a very high temperature optima assortativity coefficient (AC_t_=0.87), which quantifies the tendency of nodes being connected to similar nodes (here with similar temperature niche optima) in a network. Thus, it confirms that the latitudinal temperature gradient indirectly shaped the global ocean plankton interactome (*3*), and demonstrates the substantial effect of both environmental forcing and habitat filtering in structuring marine plankton communities at global scale.

**Fig. 1.**
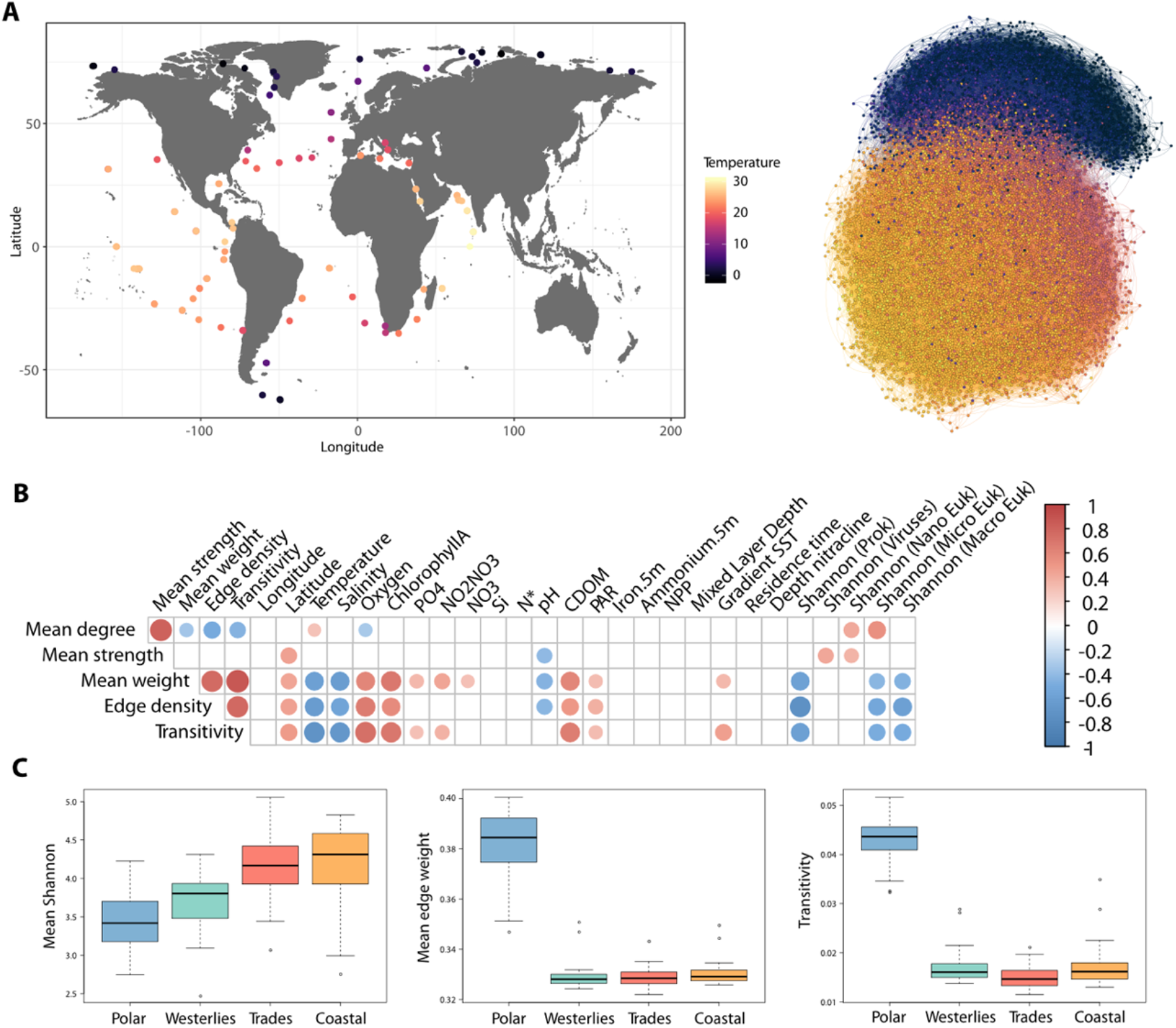
Abiotic factors shape the pole-to-pole cross-domain plankton interactome structure. (**A**) The *Tara* Oceans circumnavigation (2009-2013) included a comprehensive metabarcoding and metagenomics sampling along with physico-chemical parameters measurements covering a wide pole-to-pole latitudinal gradient of temperature. The global ocean plankton interactome (GPI) covers the three domains of life including Eukaryotes, Bacteria and Archaea and is highly structured along the latitudinal gradient of temperature from the equator to the poles. It counts 20,810 nodes (and 86,026 edges) colored according to their optimum niche temperature. (**B**) The plankton interactome topology is significantly associated to diversity, temperature, salinity, light (PAR), nutrient concentrations and pH (Spearman correlations FDR < 0.01, empty boxes correspond to non-significant correlations). (**C**) The polar interactome displays stronger associations (Mean edge weight) and clustering coefficients (transitivity) compared to other biomes (Dunn’s test, FDR < 0.05) despite its overall lower diversity.

### Abiotic factors differentially shape the plankton interactome structure

To further investigate the influence of abiotic factors in shaping the GPI structure, we extracted local subnetworks corresponding to potential interactomes at each sampling site and computed graph topological metrics. These local metrics (see supplementary text for a detailed description) were then correlated to environmental parameters (see M&M). Globally, the GPI network connectivity assessed by these metrics was negatively associated with temperature and salinity (Fig. 1B and supplementary text), pointing towards their potential impact in altering the structure of predicted interactions (*40*). We also observed a differential association between temperature and interactome connectivity in polar (Fig. S1A, negative association with strength) versus non-polar regions (positive association with strength). This difference may be linked to the observation that community turnover, which dominates in polar versus non-polar prokaryotic communities (*4*), is accompanied by stronger biotic dependencies between species. It also suggests a potential role for temperature in reducing polar community size structure in response to ocean warming, which we modeled and discuss below.

Given the observed differential association between temperature and community structure along the latitudinal axis, we compared local interactome topological metrics across biomes (Fig. 1C). The network stability (mean weight) and connectivity (transitivity) were significantly higher for the polar biome compared to other marine biomes (Dunn’s test, FDR < 0.05 for all tests), and were associated with a lower mean (cross-domains) species diversity. This higher connectivity of the polar interactome further supports the more prevalent role of biotic interactions in structuring less diverse plankton communities in the extreme polar environment. It may also reflect the lower complexity and simpler structure of pelagic ecosystems in polar oceans, which are usually characterized by shorter pathways of energy flow in food webs (*41*). The lower diversity observed in polar ecosystems has also been linked to higher prokaryotic species turnover (*42*), which may translate into the higher connectivity observed between distinct species in the polar interactome. As recently proposed for a fluvial river system, environmental heterogeneity may determine the ecological processes assembling bacterial metacommunities (*43*). Here, the environmental heterogeneity of polar systems may cause the higher network connectivity through higher heterogenous selection and community turnover.

### Biome-specific communities emerge from the plankton interactome

To further our understanding of the role of temperature in shaping the interactome structure along the latitudinal axis, we used an unsupervised approach to delineate network communities and test their association with specific biomes. Using a deterministic community detection algorithm (see M&M), five communities emerged from the GPI, which were enriched in OTUs assigned to specific biomes, and displayed distinct predicted biotic associations (Fig. 2). Through comparison of community abundance profiles, these five communities were indeed preferentially observed in specific biomes (Fig. 2A and Table S4). GPI Communities 0 and 3 (TC0 and TC3) occurred preferentially in Trades stations, Community 2 (WC2) prevailed in Westerlies stations, while Community 1 (PC1) clearly emerged in Polar stations. Community 4 (UC4) was more abundant in Polar stations but displayed a clear ubiquitous distribution. This unsupervised approach to community detection demonstrates that the GPI is self-organized across marine biomes, and that it captures the biogeography of cross-domain plankton associations. It also indicates that Longhurst’s primary biomes partitioning (*44*), which is based on chlorophyll and phenology, is also biologically meaningful for planktonic associations across plankton domains and size spectra.

**Fig. 2.**
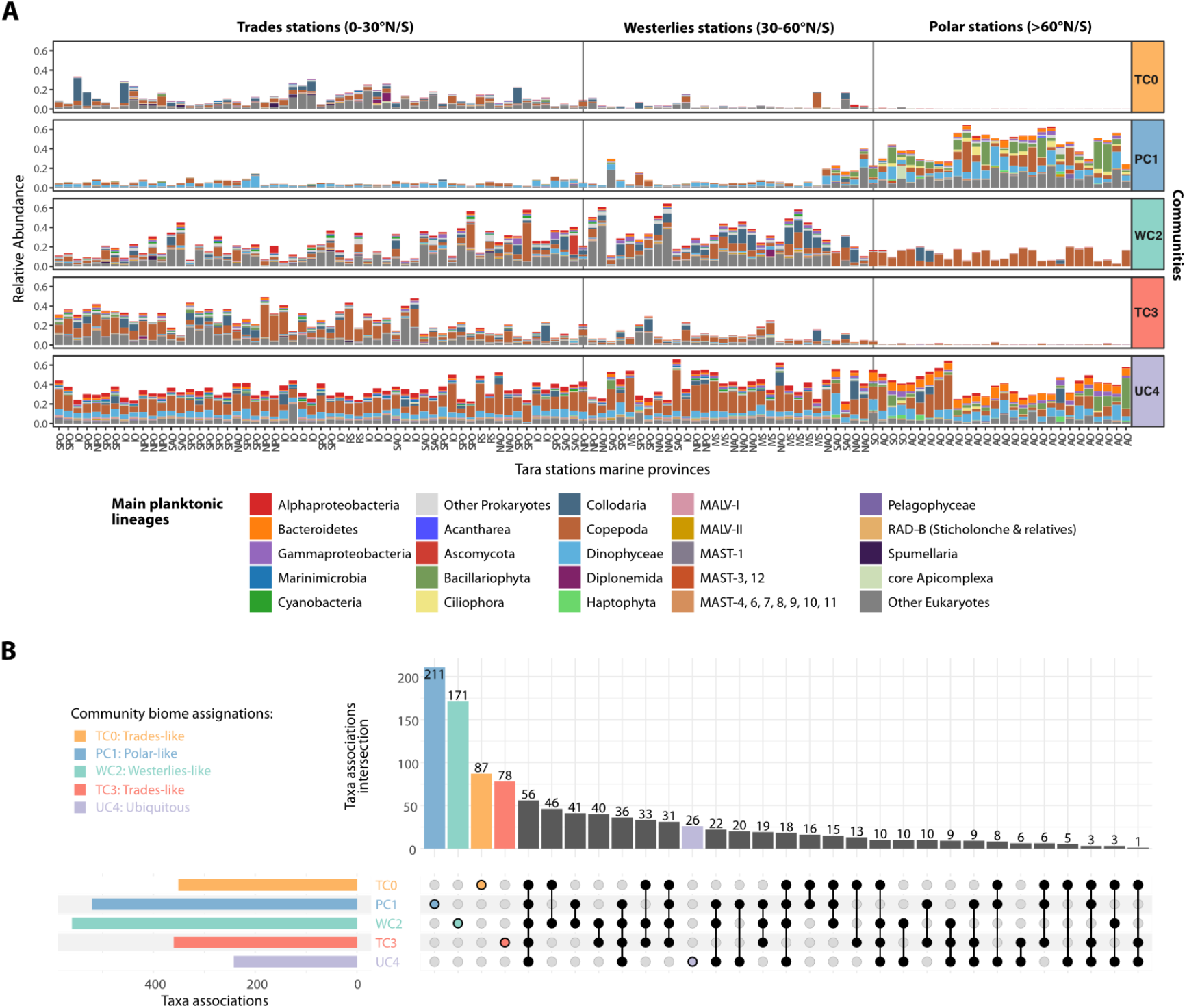
Biome-specific communities and associations emerge from the plankton interactome. (**A**) The global ocean plankton interactome (GPI) can be decomposed into 5 communities that are preferentially observed in specific marine biomes: Communities TC0 and TC3 are Trades-like, community WC2 is Westerlies-like, community PC1 is Polar-like and community UC4 is ubiquitous. Distinct main plankton lineage compositions are observed in each community along the latitudinal axis (stations are ordered by absolute latitude), disrespect of the ocean region (SPO: South Pacific Ocean, NPO: North Pacific Ocean, SAO: South Atlantic Ocean, NAO: North Atlantic Ocean, IO: Indian Ocean, RS: Red Sea, MS: Mediterranean Sea, SO: Southern Ocean, AO: Arctic Ocean). (**B**) Most plankton associations between main plankton lineages are community-specific, with communities WC2 (Westerlies-like) and PC1 (Polar-like) displaying the highest number of discriminant associations, while community UC4 displays fewer ubiquitous associations. Shared associations between communities are indicated with black filled circles and connecting lines.

All GPI communities (with the exception of the ubiquitous UC4) displayed mostly exclusive associations, even at the high taxonomic level of the main planktonic lineages considered (Fig. 2B). All GPI communities clearly differed in their associations (Fig. S3), which were enriched between distinctive taxa (Table S6). Most prevalent associations in communities TC0, TC3 and WC2 (see supplementary text) included radiolarians (e.g. Spumellaria, Acantharea and Collodaria) and Dinophyceae, detected in associations with parasitic organisms (e.g. MALV, see supplementary text for details). Both PC1 (Polar-like) and UC4 (ubiquitous) communities clearly formed two distinct systems as compared to TC0, TC3 and WC2 with respect to co-occurring lineages (Fig. S3). The PC1 community displayed a significantly lower contribution of MALV and was particularly enriched in Bacillariophyta (diatoms) associations with several eukaryotic lineages, including Ciliophora, Cryomonadida, Choanoflagellatea and Mamiellophyceae, but also with bacterial lineages such as Bacteroidetes and Gammaproteobacteria, suggesting widespread diatom-bacteria interactions (*45*) in polar ecosystems. Notably, Bacillariophyta – Cryomonadida associations may correspond to ecologically important interactions in sea-ice influenced waters. Several Cryomonadida in cold waters can feed on diatoms and some *Cryothecomonas* spp. are diatom parasitoids (*46*). The UC4 community was significantly enriched in associations involving heterotrophic bacterial lineages (Alphaproteobacteria, Gammaproteobacteria and Bacteroidetes) between themselves, and with major phytoplankton taxa such as Dinophyceae, Haptophyta, and Bacillariophyta, among the most abundant photosynthetic eukaryotes (*47*). Notable UC4 over-represented associations (Table S6) included Haptophyta with MAST and MALV lineages, emphasizing the promiscuous nature of MALV parasitic interactions, not only in tropical and temperate ecosystems (*14*), but also in polar regions. Several cross-domain associations were enriched in UC4, such as Bacillariophyta and Dinophyceae with Bacteroidetes, and Copepoda with Alphaproteobacteria, revealing the pervasive role of phytoplankton- and zooplankton-bacteria ecological interactions (*48*) shaping the plankton microbiome from pole to pole. Core associations detected across all GPI communities were also identified (n=56, Fig. 2B) and reflected strong dependencies between clades that have co-adapted to specific environmental conditions encountered in each biome. These core associations were dominated by MAST and MALV lineages, underlying their broad biogeography (*49, 50*), and very successful adaptation from pole to pole, through grazing and parasitism, respectively.

Communities emerging from the GPI underline niche differentiation by biome and implies that community-specific ecologically central species may be identified. To identify species whose impacts appear to be particularly important compared to their abundances, we computed the integrative general keystone index for each GPI community (*51*). Focusing on the ubiquitous UC4 community, the top 10 OTUs delineated by the index (Table S7) included Eukarya (n=6), among which several Copepoda (Cyclopoida, *Corycaeus* sp.) and Dinophyceae (Phalacroma, *HM581743* sp.) taxa. It also included bacterial OTUs (n=4) belonging to AEGEAN-169, NS5 marine group, *Polaribacter* and SAR116 lineages. The AEGEAN-169 group was previously shown to be abundant and ecologically important at the SPOT station (*52*). *Polaribacter* environmental genomes were recently shown to be prevalent and active in the euphotic zone at both poles (Royo-Llonch et al., submitted).

Although each GPI community was more abundant in a given biome, their occurrence goes beyond these partitions, which probably reflects the importance of physical processes (e.g. advection by ocean currents) influencing their distribution through dispersal (*53, 54*). This is also reflected by the biogeography of the WC2 community (Westerlies-like), and especially UC4 that is ubiquitous and appears to interface with other communities. The broader biogeography of these associations reflects the interconnected evolutionary history of phytoplankton- and zooplankton-bacteria ecological interactions and their pervasive role in influencing fundamental processes such as primary production, nutrient regeneration and biogeochemical cycling (*48*), not only in low-nutrient regions of the ocean but from pole to pole.

### Community-specific vulnerabilities to environmental change

Given that the GPI captured the global biogeography of cross-domain plankton associations, we sought to investigate the potential influence of environmental change on community stability across biomes. Unlike previous studies that mapped global biodiversity and investigated ecological drivers, we used the GPI as a basis to develop a novel computational framework integrating OTU niche inference and community network analyses, to assess how plankton communities and lineages may be affected under environmental change. First, for each OTU, we calculated the ecological optimum and tolerance range for a selection of environmental parameters including salinity, nutrient concentrations (NO_2_+NO_3_, PO_4_), pH and temperature. These abiotic factors are projected to change significantly under on-going climate change scenarios (*36*). For the temperature niche, we observed smaller OTU tolerance ranges towards the poles and the equator (Fig. S4), which supports the general assumption of higher environmental stability and narrower temperature ecological niches in both Polar and Trades biomes compared to the Westerlies (*44*). The environmental optima and tolerance ranges of OTUs inform us about the realized ecological niches of the taxa they represent and their potential sensitivity to environmental variations. OTUs from taxa with narrower tolerance ranges (i.e. specialists) are more likely to be affected by environmental changes, while OTUs from taxa with larger tolerance ranges (i.e. generalists) are more likely to be less sensitive to environmental changes. Based on this general assumption, we then simulated the effect of environmental changes on plankton interactome stability. Specifically, we perturbed GPI by progressively removing nodes ranked by their environmental tolerance ranges, from the narrower to the wider, for each parameter. We also attacked GPI’s nodes by their degree (i.e. from the most connected to the least connected nodes) to simulate the potentially most damaging perturbation of the network, and repeated random attacks to obtain a random expectation reference. The GPI perturbations were systematically performed at global scale, to study both global and community-specific impacts of these attacks on the stability of the network (see M&M).

In response to *in silico* environmental perturbations, we observed an overall global robustness of the network (Fig. S5). However, at local scale, we found evidence for differential effects of specific abiotic factors on GPI communities (Fig. 3A). While the UC4 (ubiquitous) community was found to be the least sensitive to simulated environmental changes (Table S8, p > 1 × 10^−2^), community TC0 (Trades-like) displayed significant vulnerabilities (Fig. S6) to temperature (Wilcoxon-rank test, p = 3.8 × 10^−10^), salinity (p = 3.8 × 10^−10^) and PO_4_ (p = 4.5 × 10^−6^), as compared to random attacks. The TC3 community (Trades-like) also displayed a significant vulnerability to temperature (p = 1.7 × 10^−3^). The WC2 community (Westerlies-like) was predicted as being the most vulnerable (Fig. S6) to nutrient concentration changes (NO_2_+NO_3_, p = 6.7 × 10^−9^; PO_4_, p = 5.2 × 10^−9^), while the PC1 community (Polar-like) displayed a clear vulnerability (Fig. 3B) to temperature (p = 3.8 × 10^−10^). The PC1 community was also found sensitive to pH (data not shown), but with lower confidence given the higher amount of missing data for pH. These distinct predicted sensitivities of GPI communities imply that taxa represented by central, most connected OTUs display lower environmental tolerance ranges for distinct abiotic factors in each community. Thus, these findings suggest that the plankton interactome will be impacted differently by environmental change in specific ecological marine regions, which are themselves predicted to be impacted differently by warming and nutrient distributions (*55*). Both Trades communities (TC0 and TC3) appeared to be more sensitive to temperature, and to a lesser extent to salinity, which are both currently increasing in tropical ocean regions (*56*). On the other hand, the Westerlies community (WC2) appeared to be more vulnerable to nutrient concentration variations, which is a coherent scenario with climate change projections (*36*). These predictions also confirm the vulnerability of the Polar community (PC1) to temperature changes that are currently occurring with the rapid warming of the Arctic over recent decades and that is projected to be amplified (*57, 58*).

**Fig. 3.**
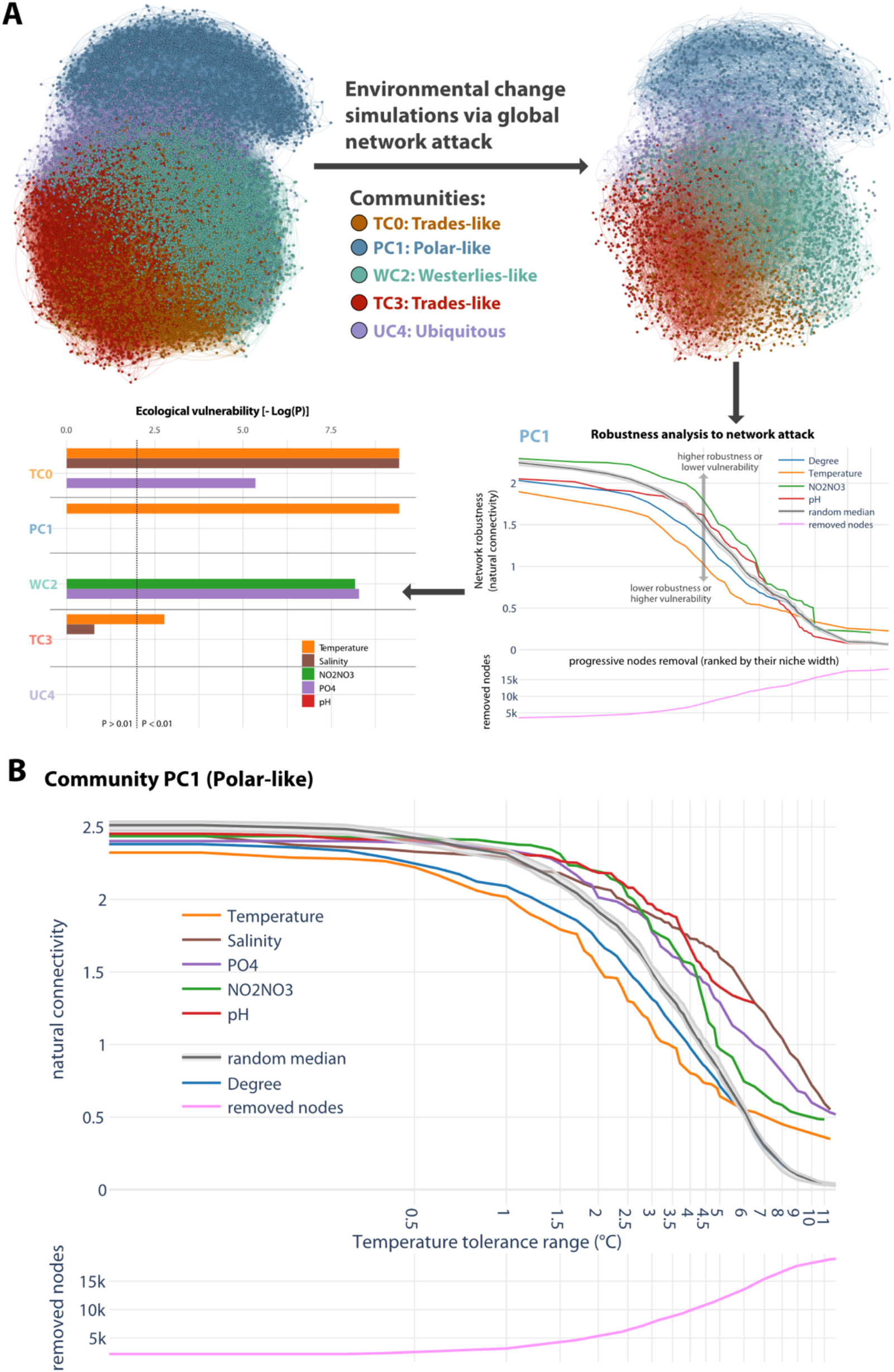
Predicting ecological vulnerabilities via network-based simulations. **(A)** Environmental change simulations are performed through tolerance range perturbations, that is progressively removing nodes of the GPI ranked by their environmental niche width (from smaller to larger), to predict ecological vulnerabilities of GPI communities. Significant vulnerabilities to environmental changes were determined by comparing distributions of the network natural connectivity (a graph robustness measure) evolution for each abiotic factor, as compared to a random perturbation. The ecological vulnerability of each GPI community was then quantified by the statistical significance (-log(P)). GPI communities TC0, TC3 (Trades-like) and PC1 (Polar) were predicted vulnerable to temperature change, while community WC2 (Westerlies-like) was predicted vulnerable to nutrient concentration variations. **(B)** The polar community (PC1) is predicted to be particularly vulnerable to temperature variations (Wilcoxon-rank test, p = 3.8 × 10^−10^).

### Plankton lineages potentially most impacted by environmental change

By combining environmental tolerance range inference with network stability analyses, plankton communities most affected by environmental perturbations were predicted, as well as vulnerabilities of the respective plankton taxa and marine plankton groups (MPGs, see M&M). For temperature vulnerability predictions, we considered relatively abundant OTUs displaying a temperature niche width smaller than 2.1 °C, which corresponds to the global mean sea surface temperature anomaly projected for the end of the century by the CMIP6 model scenario SSP2-4.5 (*59*). Marine plankton vulnerabilities to temperature, salinity and nutrient concentration changes were predicted for communities TC0 and WC2 (see supplementary text). Focusing on the PC1 polar community, which appeared the most sensitive to temperature change, we identified specific plankton lineages from all domains of life predicted to be impacted (Fig. 4A). The bacterial phyla Verrucomicrobia and Marinimicrobia were found most sensitive with a vulnerable fraction above 50%. Verrucomicrobia lineages are poorly characterized but are ubiquitous in the ocean, and may be essential for the biogeochemical cycling of carbon (*60*). Conversely, several Marinimicrobia clades have been shown to participate in the biogeochemical cycling of sulfur and nitrogen (*61*). Abundant Eukaryotic lineages for which the vulnerable fraction was above 50% included Dinophyceae, Bacillariophyta, and Ciliophora, which are all key planktonic groups in the ocean, considerably impacting global biogeochemical cycles. Notably, all MAST (Marine Stramenopiles) groups, some of which are heterotrophic and bacterivorous flagellates that interact with key photosynthetic picoplankton (*62*), are also predicted to be significantly impacted. When resuming PC1 community lineages into MPGs (Fig. 4B), we predicted a large impact from temperature changes on Archaea, phototrophs and phagotrophs, and in particular on gelatinous filter feeders. The critical role of gelatinous zooplankton within ocean trophic webs is increasingly being recognized as they may channel energy from picoplankton to higher trophic levels (*63*). The temperature sensitivity we predict for gelatinous filter feeders questions the paradigm that gelatinous zooplankton have been increasing in the past decades (*64*), and points toward the overall vulnerability of corresponding lineages to ocean warming in polar regions.

**Fig. 4.**
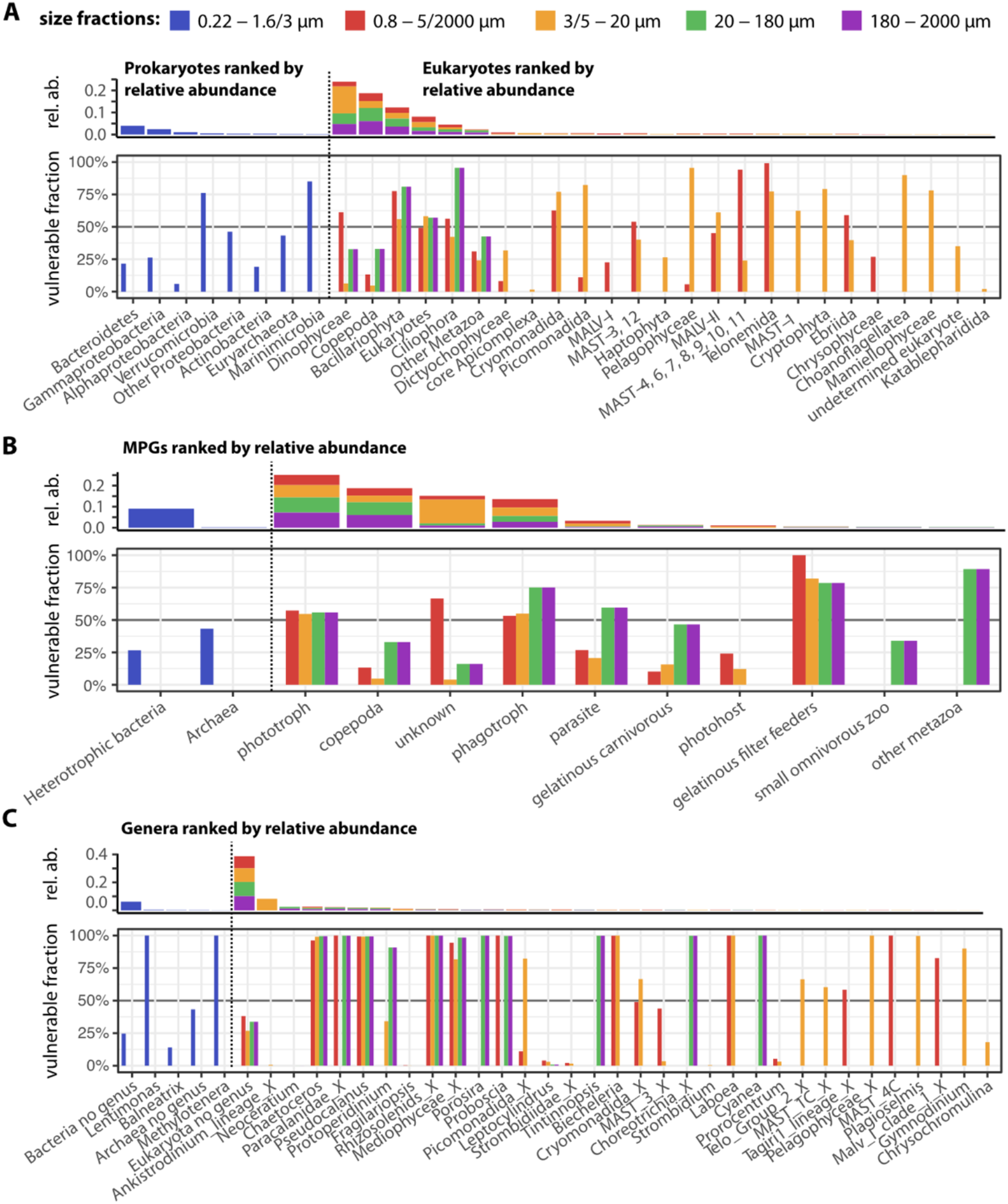
Polar marine plankton lineages and groups predicted to be most vulnerable to temperature change. (**A**) Environmental tolerance range perturbations of the GPI predicted polar marine plankton lineages (community PC1) potentially most impacted by temperature variations. **(B)** Grouping these lineages into MPGs predicted associated functions potentially most impacted by temperature variations in the polar ecosystem. **(C)** Genera most impacted by temperature variations are also identified and are potential species indicators of ocean warming in the polar ecosystem. In all panels, the fraction of lineages, MPGs, and genera (from 1 for most impacted, to 0 for not impacted) predicted to be impacted by temperature variations are depicted within each size fraction. Plankton lineages (Prokaryotes and Eukaryotes), MPGs and genera are ordered according to the cumulative mean relative abundance of the corresponding OTUs across size fractions (note that these relative abundances are not directly comparable between size fractions).

PC1 polar lineages predicted to be most sensitive to temperature were also identified at a lower taxonomic level (Fig. 4C) to infer potential species indicators of polar ecosystem change in response to ocean warming (*65*). Predicted bacterial genera as being most vulnerable to temperature change in polar regions were *Lentimonas* and *Methylotenera*, along with several uncharacterized OTUs (n=30). *Lentimonas* spp. are specialized degraders of fucoidans and other complex polysaccharides (*66*). Their observed sensitivity to temperature variations may increase the recalcitrance of algal biomass to microbial degradation, which would affect the turnover of carbon sequestered in glycans that is vital for global carbon cycling (*67*). Methylotrophs of the family Methylophilaceae play a crucial role in the carbon cycle of aquatic habitats (*68*), and several *Methylotenera* spp. isolates are methylotrophic bacteria that can use a range of one carbon compounds in coastal ocean ecosystems (*69*). Thus, these two genera appear to encompass rather specialist microbes with regard to their metabolism, and are predicted to be impacted by ocean warming in the polar ocean. Eukaryotic lineages predicted to be most sensitive to temperature included several abundant diatom genera: *Chaetoceros, Porosira* and *Proboscia*, and other genera belonging to Rhizosolenids and Mediophyceae. A single abundant genus of dinoflagellate was predicted to be impacted by temperature change: *Protoperidinium*. For copepods, the genus *Pseudocalanus* and genera from the family Paracalanidae were found to be the most vulnerable. Picomonadida was the only heterotrophic protist family predicted to be vulnerable to temperature change.

Monitoring pelagic ecosystems under environmental stress due to ongoing climate change is challenging, but plankton species indicators may provide an accurate diagnosis of ecosystem health (*70*). Previous evidence suggests that genera we predict as being most sensitive to temperature in polar ecosystems may be good candidates for plankton indicators of ocean warming. *Chaetoceros* constitutes a very large genus of marine planktonic diatoms, and is a dominant component of phytoplankton communities contributing an estimated 20% of total oceanic primary production (*71*). *Chaetoceros* is abundant in polar oceans and is affected by temperature in lab experiments (*72*). A species distribution model previously showed that the annual median probability of occurrence of another diatom species *Rhizosolenia stolterfothii* was predicted to shift in the North Atlantic Ocean (*21*), suggesting that it too will be affected by anthropogenic climate change. Considering copepods, the abundance of the genus *Pseudocalanus* has continuously decreased within a decade (2003-2012) in East Greenland waters (*73*). Another line of evidence for the temperature sensitivity of the predicted genera comes from mesocosm experiments, in which the relative biomass of a diatom from the genus *Proboscia* (*Proboscia alata*) was negatively impacted by temperature (*74*). As for dinoflagellates, a species of the genus *Protoperidinium* was shown to be less tolerant to prolonged temperature shifts in laboratory experiments (*75*).

Overall, these results underlie the differential responses of biome-specific plankton communities and associated functions to specific environmental changes. These findings provide new insights into community-specific environmental vulnerabilities of plankton lineages and associated functions. Plankton MPGs play central roles in the ecology and biogeochemistry of the polar (and global) oceans. Here, we predicted that specific plankton lineages and MPGs will be affected, which has substantial implications and may even worsen under currently projected scenarios of climate change (*59*) in the Arctic and in nutrient-rich oceanic regions.

## Conclusions

This study provides a comprehensive cross-kingdom plankton interactome covering all major oceanic provinces, including the Arctic Ocean, a region that has lacked systematic standardized sampling. Our integrated network constitutes a unique resource providing putative key ecological associations among mostly uncultivated plankton taxa identified at the molecular level. Still, this resource is limited because predicted ecological associations do not demonstrate ecological interactions (*76*), and because it does not capture the dynamics of plankton interactions that are usually assessed using temporal or longitudinal samplings (*77*). Our knowledge of plankton microbiomes, symbioses, and host-parasite relationships remains limited (*78*). While planktonic interactions remain challenging to validate, our predictions are useful to further our understanding of ecosystem functioning. Today, high-throughput co-culture experiments using microfluidics (*79*) and fabricated synthetic microbial ecosystems may help fill this gap (*80*). Our resource constitutes useful guidance for co-culture experiments, as well as a database for hypothesis testing.

Climate scenarios predict global changes in temperature, pH, and nutrient concentrations, which all greatly influence plankton physiology. Temperature can directly impact bacterial growth (*81*), grazing rates (*82*), and phytoplankton metabolism (*83*). Nitrogen availability is a primary limiting factor for marine phytoplankton (*84*). Ocean acidification caused by rising atmospheric CO_2_ can impact phytoplankton growth rates, and is predicted to have a greater impact than warming or reduced nutrient supply on plankton ecological functions (*85*). Herein, we identified and predicted distinct community vulnerabilities of the plankton interactome by studying its robustness to environmental perturbations. Overall, our findings imply differential effects of environmental change on biome-specific plankton communities resulting from biotic interactions and environmental stresses. While the influence of temperature is central, at the biome-specific community scale, salinity and nutrient concentrations were found to significantly influence plankton community structures as well. These associations support previous lines of evidence linking temperature and nutrient concentrations as the principal drivers of microbial plankton community variability (*54*).

These findings further advocate for the development of novel modeling paradigms targeting multiple biological scales (*86*), from genes to species and community levels (*87*). Our computational framework combining network analyses with niche modeling is generalizable and can be applied to various microbial ecosystems for assessing and predicting robustness to environmental perturbations. Here, specific lineage vulnerabilities were identified, but it remains an open question whether taxonomy, rather than function, is essential or not for predictive models given the potential functional redundancy in open microbial systems (*88*). Similar studies should be performed at the genomic level given that the molecular functions rather than the microbes themselves sustain marine biogeochemical processes (*89*).

## Materials and Methods

### Data description

From 2009 to 2013, the *Tara* Oceans expedition collected samples at more than 200 stations across all significant oceanic provinces from oligotrophic to polar regions. Sampling stations were selected to represent distinct marine ecosystems at global scale (*90*) describes the sampling strategy and the methodology applied, and sample provenance is described in Table S1. Environmental data measured or inferred at the depth of sampling are available in Table S2 and published at PANGAEA, Data Publisher for Earth and Environmental Science (http://www.pangaea.de). In this study, we limited our analyses to the euphotic zone, including only the samples from surface (SRF) and the Deep Chlorophyll Maximum (DCM). Two prokaryotes-enriched size fractions (0.2-1.6 µm and 0.2-3 µm) were available and included in the analyses. For Eukaryotes, the following size fractions were included (and consolidated as described below) in the analyses: ‘0.8-5 µm and 0.8-2000 µm’, ‘3-20 µm and 5-20 µm, ‘20-180 µm’ and ‘180-2000 µm. Because of these sampling constraints and the non-systematic sequencing of all available samples, the *Tara* Oceans dataset is heterogeneous. Specifically, at polar stations, fractions 0.8-5 µm and 5-20 µm are less represented. Conversely, in non-polar stations, sequencing data for the fraction 3-20 µm is nearly absent. To overcome this issue and increase sampling coverage, we considered that fractions 3-20 µm and 5-20 µm, and fractions 0.8-5 µm and 0.8-2000 µm were equivalent, as samples from these size fractions capture very similar diversity and community composition (data not shown). When both size fractions were available for the same sampling site, the 0.8-5 µm size fraction was preferred. For the 3-20/5-20 µm size fractions, only one station (TARA_124_SRF) was found to be in conflict, and we discarded the 3-20 µm sample. By doing so, we analyzed 115 sampling sites at which all considered size fractions were available.

### Data processing and taxonomic annotations

For the prokaryotes-enriched size fraction (0.2-1.6µm and 0.2-3µm), taxonomic profiling was performed using 16S ribosomal gene fragments directly identified in Illumina-sequenced metagenomes (*4*). Extracted 16S reads, named miTags, were mapped to cluster centroids of taxonomically annotated 16S rRNA gene reference sequences from the SILVA database (*59*) (release 128: SSU Ref NR 99), that had been clustered at 97% sequence identity beforehand, using USEARCH v9.2.64. Additional methodological details are available in (*4*), and we used the OTU-level abundance matrix as provided by the authors. For the Eukaryotic taxonomic profiling, we used the same methodology as in (*50*) to define OTUs using 18S rRNA gene V9 amplicons with the Swarm2 algorithm (*91*), with an updated version of the PR2 reference database (*3*). 18S rRNA gene V9 region PCR primers also amplify some 16S rRNA gene V9, thus we decided to filter out prokaryotic OTUs in the Eukaryote data that had been assigned to the following taxonomic groups: ‘Bacteria’ (35,448), ‘nan’ (31,406), ‘Archaea’ (1,806), ‘root’ (58) and ‘Organelle’ (583). For the prokaryotic abundance matrix, we filtered out miTags assigned to ‘Eukaryota’ (5,283), Chloroplast (468) and Mitochondria (74). After this filtering, we worked with six distinct matrices corresponding to each size fraction considered (see supplementary files online).

Based on these taxonomic affiliations, we classified all taxa into marine plankton groups (MPGs) as in (*3*). For prokaryotes, photosynthetic bacteria (i.e. cyanobacteria) were distinguished from heterotrophic/chemotrophic bacteria and archaea. For protists, an extended version of the functional database of (*50*) was used. It encompasses a wide variety of protist taxa that are assigned to major functional groups: photosynthetic/mixotrophic protists, endophotosymbionts, hosts with endophotosymbionts (photohosts), parasitic protists, and free-living heterotrophs or phagotrophs (heterotrophic protists). For the mesozooplankton, the categories used corresponded to the most abundant taxonomic groups (such as copepods and chaetognaths) or feeding strategies.

Shannon diversity indices were calculated for each sample and provided by (*3*).

### Data transformation and filtering

All OTU abundance matrices were transformed using the centered log-ratio (CLR) transformation (*92*). The CLR transformation is widely used in microbiome data analysis, especially in association network reconstruction (*93*), as it copes with the compositional nature of microbiome data. As log transformation cannot be applied to zero values, we added beforehand a pseudo count of one to all elements of the matrix. Finally, to reduce the high dimensionality of our data, which may be the source of false-positive predicted associations, we filtered each abundance matrix using a top-quartile filtering approach. For each sample, the upper quartile (Q3) of its non-zero abundance values was computed. An OTU was retained when its observed abundance was higher than Q3 in at least 5 samples.

### Network inference and stability procedure

The network inference was performed using FlashWeave (FW) v0.13.1 with default parameters (*38*). FW relies on the Local-to-Global learning framework and infers direct associations by searching for conditional dependencies between OTUs. Several heuristics are then applied to connect these “local” dependencies and infer a network. FW is significantly faster than other methods while achieving better or similar results, and gives the possibility to include meta-variables (such as the temperature). Although the latter feature seemed appealing, very few OTU-environmental factor associations were detected which advocates for developing a complementary approach to study the environmental influence (see Network attack below). While FW includes a heterogeneous mode (FlashWeaveHE) and the Tara dataset is heterogenous itself, the low number of samples prevented its use. Thus, we used FW in ‘sensitive’ mode without its embedded normalization since it was performed upstream to comply to our network inference strategy designed to deal with the multiple size fractions context described below.

We reconstructed graphs for each size fraction separately, running FW on the corresponding CLR-transformed abundance matrix. This first step only allows to discover *intra-fraction* edges. To connect together the 5 resulting graphs, and thus infer *inter-fraction* edges, we considered all 10 combinations of two size fractions and ran FW on the according concatenated matrices. This results in a *meta-graph* with OTUs from different size fractions being connected together.

To assess the robustness of intra- and inter-fraction edges, and reduce the number of putative false-positive associations, we implemented a *stability procedure* inspired by the STARS model selection approach (*94*). As we did for two size fractions matrices, we built every combination of three size fractions matrices and obtained 10 three-fractions graphs. We then evaluated the stability of every *meta-graph* edge by computing its frequency in the three-fractions graphs. This procedure computes a *relative stability* metric reflecting a given edge robustness to variation in both the number of samples and the number of OTUs. Edges with relative stability below 50% were removed from the meta-graph.

### Estimation of false discovery rate

Three null models were generated using two R packages (picante v1.7 and HMP v1.6). The HMP library provides the *Dirichlet*.*multinomial* function which allows data matrix generation of OTUs following a Dirichlet distribution. *picante* comes with a *randomizeMatrix* function and several methods to randomize the matrix. We used the *frequency* (that maintains OTUs occurrence frequency), and *trialswap* (maintaining OTUs occurrence frequency and sample OTUs richness) approaches. Then, networks were inferred from these matrices using FW and the same procedure as for the observed matrices. We then estimated a False Discovery Rate by comparing common edges between the observed and simulated networks. The highest FDR we obtained was 3.6% (with a number of iterations set to 10^e^8) using the *trialswap* method.

### Literature-based validation of predicted interactions

To compare the performance and sensitivity of FW to similar co-occurrence network inference methods such as SPIEC-EASI (*93*), we estimated the graph accuracy by comparing edges with known (marine) biotic interactions. We limited our comparisons to Polar networks and compared edges with known interactions from the PIDA (*78*) (https://github.com/ramalok/PIDA) and GLOBI (*95*) databases (https://www.globalbioticinteractions.org/). We used the NCBI taxonomy for prokaryotes and PR2 taxonomy for eukaryotes to identify Superkingdom, Family, Genus, and Species levels. Then, we searched for known interactions from these databases in the networks by detecting all combinations of OTUs at the four taxonomic levels considered. Results are presented in Fig. S11. Conserved associations across taxonomy ranks were estimated as follows. First, taxonomic ranks were extracted from NCBI Taxonomy database for prokaryotes and from PR2 database for eukaryotes. Next, for each pair of ranks, we counted the number of edges between nodes of each rank. Next, we repeated the procedure but now applied to the subnetwork induced by considering only nodes from a particular biome. Finally, we calculated the proportion of edges for each rank pair in each biome with respect to the total network.

### Station-specific network extraction

To further explore the association between plankton community structures and abiotic factors, we extracted sampling station-specific subnetworks corresponding to local GPI interactomes containing only nodes of OTUs detected at a given sampling station. This procedure enabled the computation of graph topological metrics (mean degree, edge density, mean weight, mean strength and transitivity) for each sampling station and enabled us to directly associate environmental parameters to local community structures.

### Marine biome assignations to OTUs

In the *Tara* Oceans data set, each sample is associated with one specific marine biome (Coastal, Trades, Westerlies or Polar). Using this information, we assigned each OTU to a biome or a combination of biomes according to its abundance profile. We did this by identifying biome(s) in which a given OTU is over-represented, based on relative abundance, compared to other biomes using a Kruskal-Wallis (KW) test implemented in the Python package SciPy (version 1.2.1). Adjustments for multiple testing were performed using the Benjamini-Hochberg (BH) procedure implemented in statsmodels (version 0.9.0). For significant tests (FDR < 0.05), a post hoc Dunn’s test implemented in scikit-posthocs (version 0.6.1) was performed to determine in which biome(s) a given OTU was significantly over-represented (FDR < 0.05). To determine the direction of the over-representation, we compared the mean values to identify and discard the “lower mean biome(s)” from the list of the OTU-associated biomes. In the GPI we were able to assign biome(s) to a significant fraction of OTUs (41.1%). Numerical and categorical assortativities were determined with the corresponding functions from networkx 2.3.

### Network community detection and biome assignation to communities

We detected five communities in the GPI using an eigenvector-based network community detection algorithm (*96*) implemented in the networkx 2.3 python package. To assign biomes to these communities, OTUs abundance tables were CLR transformed and aggregated by community for each size fraction. CLR values for each community were grouped by biome, and a KW test was run to verify mean differences of communities among biomes (KW test column in Table S4). As all p-values were significant while controlling the FDR using the BH procedure, post hoc Dunn tests were performed to detect community pairwise differences between biomes (Dunn test p-value columns in Table S4). Biomes that were found significantly lower via the Dunn test were discarded from the biome assignation (Dunn test z-score column in Table S4). The five GPI communities were found prevalent in the Polar (n=2), Westerlies (n=1) or Trades (n=2) biomes.

### Environmental optimum and tolerance range inference

Environmental optimum and tolerance range were calculated with the robust optimum (RO) method described in (*39*). For each OTU and a selection of environmental parameters, we determined the ecological optimum reflecting the optimal OTU living conditions relative to a given environmental parameter, and a tolerance range around this optimum defined by lower and upper bounds. Here, Total Sum Scaling (i.e. read count divided by the total number of reads in each sample) normalization was applied to raw matrices. For each OTU, the proportion of observed counts in a given sample is computed relatively to all samples. We use these proportions to fill a weighted vector of a fixed size (n = 10000) with environmental values accordingly (i.e. if the proportion of observed counts for OTU1 in sample 1 represents 5% of the OTU1 abundance across all samples, then the weighted vector will be filled at 5% with the environmental value measured for sample 1). The ecological optimum is then defined as the median value (Q2) of this vector, the lower and upper limits as the first (Q1) and third quartile (Q3) respectively, and the tolerance (niche) range is given by the interquartile range (Q3-Q1). Some environmental parameter values are missing (non-available, NA) for some samples. To avoid inferring spurious ecological optima and tolerance ranges for OTUs for which many values are missing, we set a minimum threshold of 10 OTU observations with non-NAs and overall with a minimum of 30% non-NAs values for it to be computed.

### General keystone index

The generalized keystone index (*51*) combines several centrality metrics in a single measure, which can then be used to rank nodes, revealing their topological importance in the network. Degree, betweenness, closeness and subgraph centralities have been calculated using the Python library networkx (version 2.3), capturing the relevance of each node at different topological scales. Factor analysis was performed with the Python library sklearn (version 0.20.3) on those centralities to get the generalized keystone index associated with each node.

### Network-based robustness analyses

In order to simulate the effects of environmental change and predict their impact on the stability of plankton community structures, we designed a network attack procedure mimicking the potential effect of each environmental parameter’s variations onto the GPI. We progressively removed network nodes by bins (n = 200 nodes until the 10000^th^ node, then n = 1000 nodes) corresponding to environmental ranges, ordered from the smallest to the largest tolerance ranges for each parameter (within a given range, the nodes are randomly sorted). At each step we computed the graph natural connectivity (*97*), a graph robustness metric, for the global interactome and for subgraphs corresponding to communities extracted from the GPI (see M&M section ‘Biome assignment and network community detection’). By doing so we could evaluate the vulnerability (or loss of robustness / stability) of the GPI at the global and community levels, and detect OTUs and lineages that were actually targeted/impacted first in the process.

Importantly, temperature and nutrient concentration changes are generally not independent; temperature increases metabolic rates, which may in turn increase nutrient uptake and cycling through the food web. Thus, both parameters may show a synergistic effect on plankton community structure (*98*). The potential for this abiotic synergy points toward a limitation of our *in silico* perturbation experiments since we did not integrate *per se* the whole set of environmental parameters that are necessary to properly define the ecological niche of a given OTU – nor the synergistic interactions between them. While an equal combination of different environmental tolerances can be assumed to define a niche (i.e. the cardinal product of limiting abiotic factors), we argue this would bias our predictions since plankton species are differentially adapted and respond to environmental conditions. Lineage-specific adaptation may explain the differential sensitivity in the Polar biome where temperature is significantly lower compared to non-polar regions, and in the Westerlies biome where nutrients are usually not limiting factors as compared to the Trades biome.

### Predicting most vulnerable community lineages and MPGs

In order to predict community-specific groups (phyla, PMGs and MPGs) most vulnerable to environmental change, we focused on most “abundant” OTUs, for which the total mean abundance was above 0.001. This cutoff corresponds to the mean relative abundance of all groups. The proportion of affected groups was computed as the factor between the total mean abundance and the affected mean abundance of a given group. For computing these affected proportions we limited ourselves to environmental ranges corresponding to global mean anomalies projected for the end of the century by the CMIP6 model scenario SSP2-4.5 (*59*). Thus, environmental ranges considered here were 2.1°C for temperature, 0.5 PSS-78 for salinity, 0.7 µmol/L for NO_2_ and 1.0 µmol/L for PO4.

### Statistical analyses

Spearman correlations, followed by BH procedure (FDR < 0.01), were performed to test associations between network topology metrics and environmental parameters (Fig. 2A). KW tests followed by Post-hoc Dunn’s tests were performed using R (version 3.2.2) to determine significant differences across biome-specific interactome topological metrics (Fig. 2C). A Pearson’s Chi-squared test was performed to detect taxa associations enriched in each interactome community (Fig. S3). Here, only pairs of taxa which co-occur in at least 3 communities and occur at minimum 50 times in total were tested. For these pairs, we performed a post hoc analysis for Pearson’s Chi-squared test on the residuals (*99*) using the chisq.posthoc.test R package (https://chisq-posthoc-test.ebbert.nrw/) to identify within each community taxa pairs with a number of associations significantly diverging from random expectation. Wilcoxon’s rank sum tests were performed to compare distributions of natural connectivity for network environmental perturbations versus random perturbations (Fig. 3).

## General

*Tara* Oceans (which includes both the *Tara* Oceans and *Tara* Oceans Polar Circle expeditions) would not exist without the leadership of the Tara Ocean Foundation and the continuous support of 23 institutes (http://oceans.taraexpeditions.org).

## Funding

We further thank the commitment of the following sponsors: CNRS (in particular Groupement de Recherche GDR3280 and the Research Federation for the study of Global Ocean Systems Ecology and Evolution, FR2022/Tara Oceans-GOSEE), European Molecular Biology Laboratory (EMBL), Genoscope/CEA, The French Ministry of Research, and the French Government ‘Investissements d’Avenir’ programmes OCEANOMICS (ANR-11-BTBR-0008), FRANCE GENOMIQUE (ANR-10-INBS-09-08), MEMO LIFE (ANR-10-LABX-54), PSL* Research University (ANR-11-IDEX-0001-02), ETH and the Helmut Horten Foundation, MEXT/JSPS/KAKENHI (projects 16H06429, 16K21723, 16H06437, 18H02279), the Spanish Ministry of Economy and Competitiveness (project MAGGY - CTM2017-87736-R), ERC Advanced Award Diatomic (grant agreement 835067 to CB), the CNRS MITI through the interdisciplinary program *Modélisation du Vivant* (GOBITMAP grant to SC), and the H2020 project AtlantECO (award number 862923). We also thank the support and commitment of Agnès b. and Etienne Bourgois, the Prince Albert II de Monaco Foundation, the Veolia Foundation, Region Bretagne, Lorient Agglomeration, Serge Ferrari, Worldcourier, and KAUST. The global sampling effort was enabled by countless scientists and crew who sampled aboard the Tara from 2009-2013, and we thank MERCATOR-CORIOLIS and ACRI-ST for providing daily satellite data during the expedition. ED is supported by the RFI ATLANSTIC2020 grant (PROBIOSTIC grant to DE). M.B. received financial support from the French Facility for Global Environment (FFEM) as part of the “Ocean Plankton, Climate and Development” project. PCJ was supported by Fundação de Amparo à Pesquisa do Estado de São Paulo – FAPESP (PhD grant 2017/26786-1). HS is supported by a Brazilian Research Council (CNPq) productivity grant (process 309514/2017-7). We are also grateful to the countries who graciously granted sampling permissions. Computational support was provided by the bioinformatics core facility of Nantes (BiRD - Biogenouest), Nantes Université, France. The authors declare that all data reported herein are fully and freely available from the date of publication, with no restrictions, and that all of the analyses, publications, and ownership of data are free from legal entanglement or restriction by the various nations whose waters the *Tara* Oceans expeditions sampled in. This article is contribution number XXX of *Tara* Oceans.

## Author contributions

S.C. designed the study. S.C., E.D., M.B., D.V. and N.H. performed the experiments. S.C., E.D., M.B., D.V. and N.H. analyzed the data. E.D. and D.V. performed the simulations. S.C., E.D. and D.E. wrote the paper, with input from M.B.S and C.B., as well as all authors.

## Competing interests

The authors state they are no competing interests.

## Data and materials availability

Data described herein are available at the EBI under the project identifiers PRJEB402 and PRJEB7988, and at PANGAEA (*90*). A web server for exploring and searching the global ocean interactome is available at: https://saas.ls2n.fr/Tara-Oceans-interactome/.

## *Tara* Oceans Coordinators (alphabetical order)

Silvia G. Acinas, Marcel Babin, Peer Bork, Emmanuel Boss, Chris Bowler, Guy Cochrane, Colomban de Vargas, Michael Follows, Gabriel Gorsky, Nigel Grimsley, Lionel Guidi, Pascal Hingamp, Daniele Iudicone, Olivier Jaillon, Stefanie Kandels-Lewis, Lee Karp-Boss, Eric Karsenti, Fabrice Not, Hiroyuki Ogata, Stéphane Pesant, Nicole Poulton, Jeroen Raes, Christian Sardet, Sabrina Speich, Lars Stemmann, Matthew B. Sullivan, Shinichi Sunagawa and Patrick Wincker.

## Notes

### Competing Interest Statement

The authors have declared no competing interest.

## References

1. C. B. Field, M. J. Behrenfeld, J. T. Randerson, P. Falkowski, Primary production of the biosphere: integrating terrestrial and oceanic components. Science 281, 237–240 (1998).

2. A. C. Gregory et al., Marine DNA Viral Macro- and Microdiversity from Pole to Pole. Cell 177, 1109–1123 e1114 (2019).

3. F. M. Ibarbalz et al., Global Trends in Marine Plankton Diversity across Kingdoms of Life. Cell 179, 1084–1097 e1021 (2019).

4. G. Salazar et al., Gene Expression Changes and Community Turnover Differentially Shape the Global Ocean Metatranscriptome. Cell 179, 1068–1083 e1021 (2019).

5. R. Stocker, Marine microbes see a sea of gradients. Science (New York, N.Y.) 338, 628–633 (2012).

6. P. G. Falkowski, R. T. Barber, V. Smetacek, Biogeochemical Controls and Feedbacks on Ocean Primary Production. 281, 200–206 (1998).

7. a. Z. Worden et al., Rethinking the marine carbon cycle: Factoring in the multifarious lifestyles of microbes. Science 347, 1257594–1257594 (2015).

8. S. Sunagawa et al., Ocean plankton. Structure and function of the global ocean microbiome. Science 348, 1261359 (2015).

9. E. Bairey, E. D. Kelsic, R. Kishony, High-order species interactions shape ecosystem diversity. Nat Commun 7, 12285 (2016).

10. D. Lawrence et al., Species interactions alter evolutionary responses to a novel environment. PLoS biology 10, e1001330 (2012).

11. J. A. Fuhrman, Microbial community structure and its functional implications. Nature 459, 193–199 (2009).

12. K. Faust, J. Raes, Microbial interactions: from networks to models. Nat Rev Microbiol 10, 538–550 (2012).

13. S. Chaffron, H. Rehrauer, J. Pernthaler, C. von Mering, A global network of coexisting microbes from environmental and whole-genome sequence data. Genome research 20, 947–959 (2010).

14. G. Lima-Mendez et al., Ocean plankton. Determinants of community structure in the global plankton interactome. Science 348, 1262073 (2015).

15. J. A. Cram et al., Cross-depth analysis of marine bacterial networks suggests downward propagation of temporal changes. ISME J 9, 2573–2586 (2015).

16. L. Guidi et al., Plankton networks driving carbon export in the oligotrophic ocean. Nature 532, 465 (2016).

17. M. T. Agler et al., Microbial Hub Taxa Link Host and Abiotic Factors to Plant Microbiome Variation. PLoS Biol 14, e1002352 (2016).

18. C. M. Herren, K. D. McMahon, Keystone taxa predict compositional change in microbial communities. Environmental Microbiology 20, 2207–2217 (2018).

19. S. Banerjee, K. Schlaeppi, M. G. A. van der Heijden, Keystone taxa as drivers of microbiome structure and functioning. Nature Reviews Microbiology 16, 567–576 (2018).

20. D. M. Needham et al., Dynamics and interactions of highly resolved marine plankton via automated high-frequency sampling. ISME J 12, 2417–2432 (2018).

21. A. D. Barton, A. J. Irwin, Z. V. Finkel, C. A. Stock, Anthropogenic climate change drives shift and shuffle in North Atlantic phytoplankton communities. Proc Natl Acad Sci U S A 113, 2964–2969 (2016).

22. D. A. Hutchins, F. Fu, Microorganisms and ocean global change. Nat Microbiol 2, 17058 (2017).

23. G. Beaugrand, R. R. Kirby, How Do Marine Pelagic Species Respond to Climate Change? Theories and Observations. Annual Review of Marine Science 10, 169–197 (2018).

24. G. Beaugrand et al., Prediction of unprecedented biological shifts in the global ocean. Nature Climate Change 9, 237–243 (2019).

25. J. A. Wiens, D. Stralberg, D. Jongsomjit, C. A. Howell, M. A. Snyder, Niches, models, and climate change: assessing the assumptions and uncertainties. Proc Natl Acad Sci U S A 106 Suppl 2, 19729–19736 (2009).

26. P. Frémont et al., Restructuring of genomic provinces of surface ocean plankton under climate change. bioRxiv, (2020).

27. A. Guisan et al., Making better biogeographical predictions of species’ distributions. Journal of Applied Ecology 43, 386–392 (2006).

28. J. A. Fuhrman, J. A. Cram, D. M. Needham, Marine microbial community dynamics and their ecological interpretation. Nat Rev Microbiol 13, 133–146 (2015).

29. C. Moore, J. Grewar, G. S. Cumming, Quantifying network resilience: comparison before and after a major perturbation shows strengths and limitations of network metrics. Journal of Applied Ecology 53, 636–645 (2016).

30. M. B. Araújo, M. Luoto, The importance of biotic interactions for modelling species distributions under climate change. Global Ecology and Biogeography 16, 743–753 (2007).

31. G. E. P. Murphy, T. N. Romanuk, B. Worm, Cascading effects of climate change on plankton community structure. Ecology and Evolution 10, 2170–2181 (2020).

32. R. V. Sole, J. M. Montoya, Complexity and fragility in ecological networks. Proc Biol Sci 268, 2039–2045 (2001).

33. G. E. G. E. Hutchinson, An introduction to population ecology. (Yale University Press, 1978).

34. P. Brun et al., Ecological niches of open ocean phytoplankton taxa. Limnology and Oceanography 60, 1020–1038 (2015).

35. C.L. Quéré et al., Ecosystem dynamics based on plankton functional types for global ocean biogeochemistry models. Global Change Biology 11, 2016–2040 (2005).

36. L. Bopp et al., Multiple stressors of ocean ecosystems in the 21st century: projections with CMIP5 models. Biogeosciences 10, 6225–6245 (2013).

37. E. Karsenti et al., A holistic approach to marine eco-systems biology. PLoS biology 9, e1001177 (2011).

38. J. Tackmann, J. F. Matias Rodrigues, C. von Mering, Rapid inference of direct interactions in large-scale ecological networks from heterogeneous microbial sequencing data. bioRxiv, 390195 (2018).

39. E. Cristóbal, S. V. Ayuso, A. Justel, M. Toro, Robust optima and tolerance ranges of biological indicators: a new method to identify sentinels of global warming. Ecological Research 29, 55–68 (2014).

40. M. Dossena et al., Warming alters community size structure and ecosystem functioning. Proc Biol Sci 279, 3011–3019 (2012).

41. E. J. Murphy et al., Understanding the structure and functioning of polar pelagic ecosystems to predict the impacts of change. Proc Biol Sci 283, (2016).

42. D. Righetti, M. Vogt, N. Gruber, A. Psomas, N. E. Zimmermann, Global pattern of phytoplankton diversity driven by temperature and environmental variability. Sci Adv 5, eaau6253 (2019).

43. P. Huber et al., Environmental heterogeneity determines the ecological processes that govern bacterial metacommunity assembly in a floodplain river system. ISME J, (2020).

44. A. R. Longhurst, in Ecological Geography of the Sea (Second Edition), A. R. Longhurst, Ed. (Academic Press, Burlington, 2007), pp. 89–102.

45. S. A. Amin, M. S. Parker, E. V. Armbrust, Interactions between diatoms and bacteria. Microbiol Mol Biol Rev 76, 667–684 (2012).

46. M. Thaler, C. Lovejoy, Distribution and diversity of a protist predator Cryothecomonas (Cercozoa) in Arctic marine waters. J Eukaryot Microbiol 59, 291–299 (2012).

47. J. J. Pierella Karlusich, F. M. Ibarbalz, C. Bowler, Phytoplankton in the Tara Ocean. Ann Rev Mar Sci 12, 233–265 (2020).

48. J. R. Seymour, S. A. Amin, J. B. Raina, R. Stocker, Zooming in on the phycosphere: the ecological interface for phytoplankton-bacteria relationships. Nat Microbiol 2, 17065 (2017).

49. Y. Seeleuthner et al., Single-cell genomics of multiple uncultured stramenopiles reveals underestimated functional diversity across oceans. Nat Commun 9, 310 (2018).

50. C. de Vargas et al., Ocean plankton. Eukaryotic plankton diversity in the sunlit ocean. Science 348, 1261605 (2015).

51. E. Estrada, Characterization of topological keystone species: Local, global and “meso-scale” centralities in food webs. Ecological Complexity 4, 48–57 (2007).

52. J. A. Cram et al., Seasonal and interannual variability of the marine bacterioplankton community throughout the water column over ten years. ISME J 9, 563–580 (2015).

53. D. J. Richter et al., Genomic evidence for global ocean plankton biogeography shaped by large-scale current systems. bioRxiv, (2019).

54. R. Logares et al., Disentangling the mechanisms shaping the surface ocean microbiota. Microbiome 8, 55 (2020).

55. J. K. Moore et al., Sustained climate warming drives declining marine biological productivity. Science 359, 1139–1143 (2018).

56. P. J. Durack, S. E. Wijffels, R. J. Matear, Ocean salinities reveal strong global water cycle intensification during 1950 to 2000. Science 336, 455–458 (2012).

57. T. G. Shepherd, Effects of a warming Arctic. Science 353, 989–990 (2016).

58. E. Post et al., The polar regions in a 2 degrees C warmer world. Sci Adv 5, eaaw9883 (2019).

59. L. Kwiatkowski et al., Twenty-first century ocean warming, acidification, deoxygenation, and upper-ocean nutrient and primary production decline from CMIP6 model projections. Biogeosciences 17, 3439–3470 (2020).

60. S. Freitas et al., Global distribution and diversity of marine Verrucomicrobia. ISME J 6, 1499–1505 (2012).

61. A. K. Hawley et al., Diverse Marinimicrobia bacteria may mediate coupled biogeochemical cycles along eco-thermodynamic gradients. Nat Commun 8, 1507 (2017).

62. Y. C. Lin et al., Distribution patterns and phylogeny of marine stramenopiles in the north pacific ocean. Appl Environ Microbiol 78, 3387–3399 (2012).

63. C. Jaspers, J. L. Acuña, R. D. Brodeur, Interactions of gelatinous zooplankton within marine food webs. Journal of Plankton Research 37, 985–988 (2015).

64. R. H. Condon et al., Questioning the Rise of Gelatinous Zooplankton in the World’s Oceans. BioScience 62, 160–169 (2012).

65. D. Kalenitchenko, N. Joli, M. Potvin, J.-É. Tremblay, C. Lovejoy, Biodiversity and Species Change in the Arctic Ocean: A View Through the Lens of Nares Strait. Frontiers in Marine Science 6, (2019).

66. A. Sichert et al., Verrucomicrobia use hundreds of enzymes to digest the algal polysaccharide fucoidan. Nat Microbiol 5, 1026–1039 (2020).

67. F. Cuskin, E. C. Lowe, Glycan degradation writ large in the ocean. Nat Microbiol 5, 980–981 (2020).

68. L. Chistoserdova, Methylotrophs in natural habitats: current insights through metagenomics. Appl Microbiol Biotechnol 99, 5763–5779 (2015).

69. A. Ramachandran, D. A. Walsh, Investigation of XoxF methanol dehydrogenases reveals new methylotrophic bacteria in pelagic marine and freshwater ecosystems. FEMS Microbiol Ecol 91, (2015).

70. G. Beaugrand, Monitoring pelagic ecosystems using plankton indicators. ICES Journal of Marine Science 62, 333–338 (2005).

71. S. Malviya et al., Insights into global diatom distribution and diversity in the world’s ocean. Proceedings of the National Academy of Sciences of the United States of America 113, E1516–E1525 (2016).

72. X. Li, N. Roevros, F. Dehairs, L. Chou, Biological responses of the marine diatom Chaetoceros socialis to changing environmental conditions: A laboratory experiment. PLoS One 12, e0188615 (2017).

73. K. E. Arendt, M. D. Agersted, M. K. Sejr, T. Juul-Pedersen, Glacial meltwater influences on plankton community structure and the importance of top-down control (of primary production) in a NE Greenland fjord. Estuarine, Coastal and Shelf Science 183, 123–135 (2016).

74. U. Sommer, C. Paul, M. Moustaka-Gouni, Warming and Ocean Acidification Effects on Phytoplankton--From Species Shifts to Size Shifts within Species in a Mesocosm Experiment. PLoS One 10, e0125239 (2015).

75. G. Franzè, S. Menden-Deuer, Common temperature-growth dependency and acclimation response in three herbivorous protists. Marine Ecology Progress Series 634, 1–13 (2020).

76. F. G. Blanchet, K. Cazelles, D. Gravel, Co-occurrence is not evidence of ecological interactions. Ecology Letters 23, 1050–1063 (2020).

77. J. A. Gilbert et al., Defining seasonal marine microbial community dynamics. ISME J 6, 298–308 (2012).

78. M. F. M. Bjorbaekmo, A. Evenstad, L. L. Rosaeg, A. K. Krabberod, R. Logares, The planktonic protist interactome: where do we stand after a century of research? ISME J 14, 544–559 (2020).

79. M. Girault, T. Beneyton, Y. Del Amo, J. C. Baret, Microfluidic technology for plankton research. Curr Opin Biotechnol 55, 134–150 (2019).

80. K. Zengler et al., EcoFABs: advancing microbiome science through standardized fabricated ecosystems. Nat Methods 16, 567–571 (2019).

81. D. L. Kirchman, X. A. Moran, H. Ducklow, Microbial growth in the polar oceans - role of temperature and potential impact of climate change. Nat Rev Microbiol 7, 451–459 (2009).

82. H. Sarmento, J. M. Montoya, E. Vazquez-Dominguez, D. Vaque, J. M. Gasol, Warming effects on marine microbial food web processes: how far can we go when it comes to predictions? Philos Trans R Soc Lond B Biol Sci 365, 2137–2149 (2010).

83. G. Yvon-Durocher, J. I. Jones, M. Trimmer, G. Woodward, J. M. Montoya, Warming alters the metabolic balance of ecosystems. Philos Trans R Soc Lond B Biol Sci 365, 2117–2126 (2010).

84. S. Basu, R. K. Mackey, Phytoplankton as Key Mediators of the Biological Carbon Pump: Their Responses to a Changing Climate. Sustainability 10, (2018).

85. S. Dutkiewicz et al., Impact of ocean acidification on the structure of future phytoplankton communities. Nature Climate Change 5, 1002–1006 (2015).

86. M. Kumar, B. Ji, K. Zengler, J. Nielsen, Modelling approaches for studying the microbiome. Nat Microbiol 4, 1253–1267 (2019).

87. D. D’Alelio et al., Modelling the complexity of plankton communities exploiting omics potential: From present challenges to an integrative pipeline. Current Opinion in Systems Biology 13, 68–74 (2019).

88. S. Louca et al., Function and functional redundancy in microbial systems. Nat Ecol Evol 2, 936–943 (2018).

89. V. J. Coles et al., Ocean biogeochemistry modeled with emergent trait-based genomics. Science 358, 1149–1154 (2017).

90. S. Pesant et al., Open science resources for the discovery and analysis of Tara Oceans data. Sci Data 2, 150023 (2015).

91. F. Mahé, T. Rognes, C. Quince, C. de Vargas, M. Dunthorn, Swarm: robust and fast clustering method for amplicon-based studies. PeerJ 2, e593 (2014).

92. J. Aitchison, The Statistical Analysis of Compositional Data. Journal of the Royal Statistical Society. Series B (Methodological) 44, 139–177 (1982).

93. Z. D. Kurtz et al., Sparse and compositionally robust inference of microbial ecological networks. PLoS Comput Biol, 1–25 (2014).

94. H. Liu, K. Roeder, L. Wasserman, paper presented at the Proceedings of the 23rd International Conference on Neural Information Processing Systems - Volume 2, Vancouver, British Columbia, Canada, 2010.

95. J. H. Poelen, J. D. Simons, C. J. Mungall, Global biotic interactions: An open infrastructure to share and analyze species-interaction datasets. Ecological Informatics 24, 148–159 (2014).

96. M. E. Newman, Modularity and community structure in networks. Proc Natl Acad Sci U S A 103, 8577–8582 (2006).

97. W. Jun, M. Barahona, T. Yue-Jin, D. Hong-Zhong, Natural Connectivity of Complex Networks. Chinese Physics Letters 27, 078902 (2010).

98. E. Maranon, M. P. Lorenzo, P. Cermeno, B. Mourino-Carballido, Nutrient limitation suppresses the temperature dependence of phytoplankton metabolic rates. ISME J 12, 1836–1845 (2018).

99. T. M. Beasley, R. E. Schumacker, Multiple Regression Approach to Analyzing Contingency Tables: Post Hoc and Planned Comparison Procedures. The Journal of Experimental Education 64, 79–93 (1995).

