## Supplementary information for "Environmental vulnerability of the global ocean plankton community interactome"

### Supplementary text

#### Network inference

We integrated data from 575 samples derived from several size-fractions (Table S1) in the euphotic zone across nine oceanic provinces and four marine biomes as defined by Longhurst (*1*). The plankton samples included organisms from Bacteria and Archaea to pico-, nano-, micro- and macro-Eukaryotic species. We derived species abundance profiles from 16S rRNA gene reads extracted from metagenomes (miTags), and 18SV9 rRNA gene sequences for prokaryotes and eukaryotes, respectively (*2*) (Table S1). Environmental data (Table S2) were also collected from on-site and satellite measurements, as well as from the World Ocean Database (*3*).
The Tara Oceans heterogeneous sampling and the number of available samples can directly influence the topology of the network (*4*). In addition, the methods used to cluster sequences were different across datasets. 18S Operational Taxonomic Units (OTUs) were identified by clustering V9 amplicons using Swarm2 (*5*), while 16S miTags OTUs were identified using miTags reads mapped on SILVA database (*6*) reference OTUs defined at 97% identity. This heterogeneity leads to differences in terms of resolution and number of OTUs corresponding to each domain. The size fractions heterogeneity and the OTU clustering are potential biases that have to be considered when inferring co-occurrence networks. Thus, the GPI integrates only sampling stations at which similar size fractions were available, thereby avoiding this methodological drawback. Viral size fraction samples were not included to obtain a homogeneous sampling while maximizing latitudinal coverage. Co-occurrence network inference is also sensitive to the number of OTUs detected, which can impact the resulting network topology. This bias is not easily correctable as the difference can correspond to true biological diversity or may be due to distinct methods used to cluster sequences. Here, we did not correct or adjust for the number of OTUs, but we analyzed networks considering eukaryotes as probably overrepresented (due to the higher number of Eukaryotic size fractions and the OTU definition method). In addition, we generated a merged interactome in which Eukaryotic OTUs identified in more than one size fraction are merged, in this network a given OTU corresponds to a single node. This network was used for taxa and associations enrichment analyses.

Including environmental factors into the network inference identified very few edges (n = 325, 0.53%) between OTUs and environmental factors. As previously reported (*7*), this suggests that abiotic factors are incomplete predictors of plankton community structure and highlights the role of top-down biotic interactions in the epipelagic ocean. While biotic interactions have a crucial role in shaping plankton community structure, the impact of abiotic factors may be underestimated since plankton species niches are not explicitly considered when reconstructing networks. To alleviate this shortcoming and revisit the role of abiotic factors in shaping plankton community structure, we opted to exclude the abiotic factors from the network inference, but rather estimated robust ecological optima and tolerance ranges (*8*) for each OTU and available environmental parameters (see M&M and Table S3). The overall workflow we developed to infer, validate and analyze the GPI is presented in Fig. S10.

**Network validation**

Different null models were generated for validation purposes, by generating different null models we estimated a false discovery rate below 3.6% in the GPI (see M&M). We also used databases of known ecological interactions (*9, 10*) to validate polar-specific associations inferred using the current state-of-the-art methods (FlashWeave (*11*) and SPIEC-EASI (*12*)) and showed that they both performed well and recovered similar numbers of known interactions (*9*) (Fig. S11). To further validate and estimate the GPI potential to predict novel interactions, we compared its potential to recover known interactions, as in a previous interactome (*7*). The GPI doubled the number of recovered known interactions (at genus level) from the PIDA (*9*) database (n = 58 in GPI vs. n = 25) while maintaining a false discovery rate below 10% (5.5% in GPI vs. 5.6%, as estimated by considering known interactions). The GPI also doubled the number of recovered known interactions (at genus level) from the GLOBI (*10*) database (n = 110 in GPI vs. n = 52) while reducing the false discovery rate from 4.6% to 2.6%. In addition, we searched the GPI for highly specific known host-symbiont interactions (parasite and photosymbionts). Notably, it captured the specific symbiosis between *Tiarina* and *Symbiodinium* (*13*) and also the interaction between *Symsagittifera* and *Tetraselmis* (*7*), all at the very specific level of amplicon reads forming these OTUs of interest. These results provide a high confidence level in the ability of GPI to predict potential novel biotic interactions.

**Network topological metrics**

A number of topological metrics were computed for the GPI global and local station-specific networks, there are listed below with a short definition and a description of what they capture:

- Assortativity: It quantifies the tendency of nodes being connected to similar nodes in a network with respect to a given attribute (numeric or categorical).
- Mean degree: The mean number of connections or edges a given node has to other nodes, it is closely related to the density of a network.
- Mean weight: The mean of weights assigned to edges, capturing the overall strength or "stability" of the associations / predicted interactions.
- Mean strength (or weighted vertex degree): The mean sum of edge weights of the adjacent edges for each vertex.
- Edge density: The ratio between the number of edges and the number of possible edges.
- Transitivity (or clustering coefficient): A measure of the degree to which nodes in a graph tend to cluster together (community clustering).
- Natural connectivity: A measure of the redundancy of alternative paths in a network based on evaluating the weighted number of closed walks, it quantifies the robustness of a network.
- Randic's index (or connectivity index): A degree-based topological index measuring the process of connecting various parts of a network.

**Abiotic factors differentially shape the plankton interactome structure**

At global scale, temperature, salinity, light (measured as photosynthetically active radiation - PAR), nutrient concentration ($PO_4$, $NO_2$+$NO_3$, $NO_3$) and pH were significantly associated to interactome topological metrics (Fig. S1). Temperature was

negatively associated to transitivity (or clustering coefficient), which measures the tendency of nodes to cluster together, and thus supports the role of temperature in controlling the interactome structure. Similarly, salinity was negatively associated to transitivity, while associations with light, nutrient concentrations and pH were weaker. Interestingly, we also observed a positive association between transitivity and Gradient Sea Surface Temperature (Fig. S1A), a good proxy for physical processes such as ocean fronts and potential vertical transport, suggesting their role in influencing the interactome structure as well. In addition, the relationship between interactome mean strength (or weighted node degree) and temperature is noteworthy, and the observed slop inversion suggests a differential role for temperature in shaping interactome structure in polar versus non-polar systems.
Given the latter observation and the very distinct oceanographic and phenological features of polar systems, we explored the potential influence of abiotic factors on the GPI structure in polar vs. non-polar stations (Fig. S1B). In non-polar stations (Fig. S1B upper panel), transitivity remained negatively associated with temperature and salinity, confirming their putative influence in shaping predicted interactions in temperate and tropical ecosystems. The non-polar mean weight and transitivity were positively associated with chlorophyll-*a*, $NO_2$+$NO_3$, colored dissolved organic matter (CDOM) and net primary productivity (NPP) (Fig. S1C left panel) suggesting a critical role of predicted biotic interactions in modulating marine productivity (*14*). Here, temperature, salinity and diversity seemed to increase the potential for interactions (strength) (*15*). However, temperature and salinity also appeared to affect transitivity (or community clustering, Fig. S1C left panel), highlighting the combined effect of biotic and abiotic constraints in shaping non-polar plankton communities (*16*).

In the polar biome (Fig. S1B lower panel), mostly represented by Arctic stations (n = 20) rather than Southern Ocean stations (n = 3), patterns of association between abiotic factors and the GPI topology were different, suggesting that distinct physical, biological and ecological processes underpin ecosystem structuring at the poles. Here, all topological metrics were negatively associated with salinity and positively with CDOM, likely underlying the critical influence of freshwater inputs into the Arctic Ocean (*17*). Notably, both network strength and weight were linked to nitrogen cycling through nitrate and ammonium concentrations (Fig. S1C right panel), underlying the role of nitrogen limitation in shaping polar community structures, at least during summer time (*18*). Thus, salinity and nitrate ($NO_3$) concentration emerged as the main drivers forcing plankton ecological associations in the polar system. Here, the link with salinity and CDOM likely reflects the critical influence of freshwater inputs (via rivers) into the Arctic Ocean ecosystem (*19*). In contrast, the association with nitrate concentration may be linked to plankton phenology (high concentration pre-bloom/bloom, and low concentration post-bloom) that considerably impacts community composition (*20*), or the influence of N* in shaping phytoplankton ecological niches in the coastal Arctic Ocean (*21*).

**Biome-specific communities emerge from the plankton interactome**
To complement our network-based analysis and confirm the role of temperature in shaping the interactome structure along the latitudinal axis, we assigned preferential marine biomes as defined by Longhurst (Trades, Westerlies, Polar, and Coastal) to OTUs (see M&M). The GPI was found to be well structured according to marine biomes with a positive assortativity coefficient ($AC_b$=0.35).

To identify community-specific taxonomic groups, we summarized the GPI in 36 main lineages (29 Eukarya, 10 Bacteria and 3 Archaea) and performed a correspondence analysis (Fig. S2 and Table S5). This revealed the community-level biome endemicity of certain taxa in predicted associations (Fig. 2A). Considering all associations within GPI, the main planktonic lineages were highly interconnected (Fig. S3). In agreement with the first *Tara* Oceans interactome (*7*), a significant number of associations involved the marine parasitic syndiniales (MALV), dinoflagellates and arthropods, the latter mainly represented by Copepoda. The exceptions were for specific groups of bacteria (Cyanobacteria, Candidate phylum *Marinimicrobia* (formerly SAR406) and Deltaproteobacteria) that were primarily predicted to interact with other prokaryotic groups.

The GPI biome-specific communities TC0, TC3 (Trades-like) and WC2 (Westerlies-like) shared the highest numbers of taxa in associations. While TC0 and TC3 were both enriched in Collodaria, MALV lineages were more prevalent in TC0 and WC2, and Dinophyceae were more prevalent in TC3 (Fig. S2B). Communities TC0 and TC3 were poorly enriched in Bacillariophyta (diatom) associations, consistent with their higher abundance in polar and coastal environments (*22*). Community TC0 (Trades-like) displayed distinct prevalent lineages, including Spumellaria, RAD-A, Nassellaria and Acantharea. Communities PC1 (Polar-like) and UC4 (ubiquitous) were the most dissimilar and displayed very distinct prevalent lineages. Phototrophic groups such as Mamiellophyceae and Bacillariophyta, but also heterotrophic groups like filosan (Cryomonadida and other filosans), Ciliophora and MAST-1 were clearly enriched in PC1. Notably, bacterial lineages (Alphaproteobacteria, Bacteroidetes, Gammaprotebacteria and *Marinimicrobia*) and Haptophyta were particularly prevalent within the UC4 (ubiquitous) community, consistent with their high diversity and relatively low dispersal limitation (*23*). The identification of specific lineages enriched in each community underlines the strong role of temperature, habitat filtering and environmental stability in structuring marine plankton communities from pole to pole (*24*).

The most prevalent associations in the TC0 community included several radiolarians (Spumellaria, Acantharea and Collodaria), notably detected in associations with parasitic syndiniales (MALV), supporting the hypothesis that radiolarians are a reservoir for MALV (*25*). Conversely, the TC3 community was dominated by associations between Dinophyceae and MALV-II, and Collodaria and Bacillariophyta. Given that one of the few described genera of MALV-II is *Amoebophrya*, which is a parasitoid of a wide range of Dinophyceae in coastal and open waters (*26*), these predicted interactions are likely host-parasitic. The WC2 community is characterized by associations between MALV lineages and several abundant and widespread groups including Collodaria, Ciliophora, Copepoda and Dinophyceae. Collodaria is a radiolarian lineage usually found in subtropical and tropical waters that display colonial lifestyle and no silicification. Notably, Collodaria have acquired phototrophic protists that bear photosynthetic endosymbionts, providing them with fixed carbon. Here, Collodaria appeared highly connected with Dinophyceae in WC2, which supports the observation that most described marine protist endosymbionts are dinoflagellates (*27*).

**Community-specific vulnerabilities to environmental change**

The GPI network-based perturbations (via nodes removal) were defined by the environmental tolerance ranges of each OTU but did not integrate the potential effect of perturbed or lost biotic relationships. To integrate predicted plankton interactions, we also simulated environmental perturbations onto GPI by targeting edges (instead of nodes), progressively removing edges linking nodes from the smallest to the largest tolerance range (or niche) overlap (Fig. S7). This analysis assumed that all predicted GPI interactions are essential (which is not a priori true), but integrated the potential effects of both abiotic and biotic factors. This complementary simulation supported our observations of temperature sensitivity in both TC0 (Trades-like) and PC1 (Polar-like) communities, as well as the sensitivity to nutrient change ($NO_2$+$NO_3$) in the WC2 community (Westerlies-like).

**Community lineages potentially most impacted by environmental change**
Considering the salinity vulnerability of the TC0 (Trades-like) community, mainly Eukaryotic plankton lineages were predicted to be impacted, in particular members of the Collodaria, Diplonemida and Mamiellopyceae (Fig. S8). These predictions were also linked to community TC0 MPGs predicted to be most impacted by salinity variations. Here, mainly phagotrops taxa appeared most impacted by salinity change. The extreme diversity and large biogeography of Diplonemida and Collodaria have been witnessed thanks to environmental sequencing via metabarcoding (*28, 29*). Representatives of these groups are difficult to cultivate, thus constraining lab experiments. Here, we predicted that both these groups may actually be impacted by salinity changes in the tropical ocean.
Within the WC2 (Westerlies-like) community, plankton lineages predicted to be most impacted by nitrate variations were distinct, and included several Bacteria and Archaea (Fig. S9). Gammaproteobacteria, Bacteroidetes, Verrucomicrobia and Planctomycetes appeared most impacted by nitrate changes. These groups are mainly heterotrophic lineages and their vulnerabilities may have consequences for nutrients cycling in temperate oceans, notably for nitrogen fixation that has recently been shown to be carried by diazotrophic planctomycetes in the surface ocean (*30*). Most abundant eukaryotic lineages predicted to be impacted by nitrate variations included copepods, Collodaria, Dinophyceae and MALV lineages, as well as Ascomycota taxa. When summarizing WC2 community lineages into MPGs, a larger impact of nutrient concentration changes was predicted on heterotrophic bacteria, copepods and photohosts, but also on other metazoans and smaller omnivorous zooplankton. Overall, plankton groups predicted most impacted by environmental changes in WC2 included Copepods and Collodaria, two globally dominant plankton groups and major drivers of carbon export in the ocean (*14, 31*). Members of the Stramenopiles, which are ecologically important group of marine plankton (*32*), as well as several MALV lineages, which are very diverse and abundant groups in the global ocean (*33*), were also predicted vulnerable to nitrate variations.

### <u>Supplementary tables</u>

Table S1. Samples description.
Table S2. Environmental data associated with samples.
Table S3. OTU ecological optimum and tolerance range in GPI.
Table S4. Biome assignation to GPI communities.
Table S5. Over/under-represented taxa in GPI communities.
Table S6. Over/under-represented taxa associations in GPI communities.
Table S7. Keystone general indices in GPI.
Table S8. GPI community vulnerabilities to environmental parameters.

**Supplementary files**

Size-fractions filtered abundance matrices used for inferring GPI: *GPI-matrices.zip*
The GPI interactome as a graphML file: *GPI.graphml*
The GPI interactome, merged at the OTU level, as a graphML file: *GPI-merged.graphml*

A web server for exploring and searching the GPI is available at: https://saas.ls2n.fr/Tara-Oceans-interactome/.

### Supplementary figures

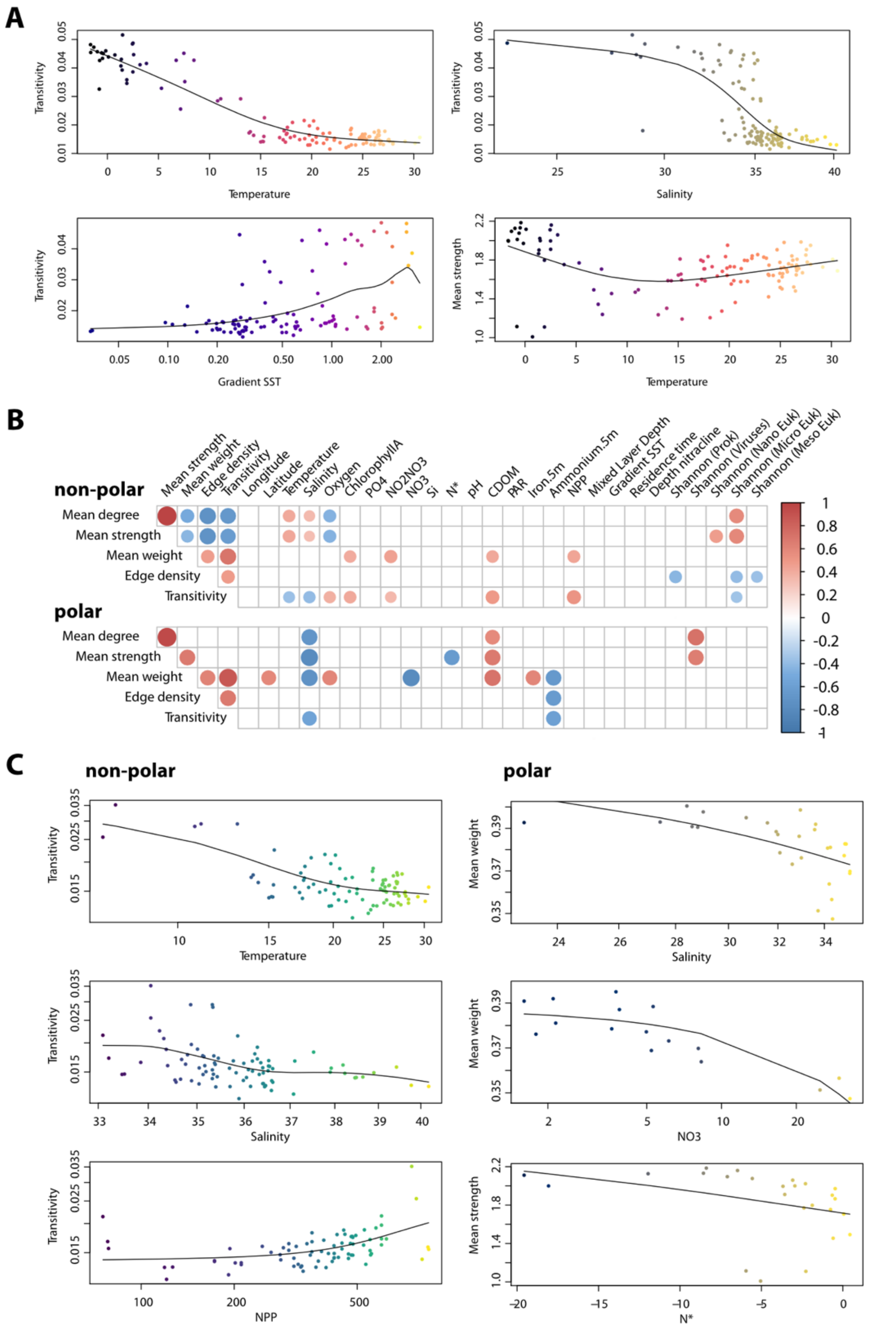


**Fig. S1. Abiotic factors differentially shape plankton interactome structure.** (**A**) Temperature and salinity emerged as the strongest drivers shaping interactome connectivity (transitivity) at global scale. The interactome transitivity is also significantly associated to gradient SST. Temperature is differentially associated to interactome strength (or weighted node degree) suggesting a differential role for temperature in shaping interactome structure in polar versus non-polar systems. (**B**) The plankton interactome topology is significantly associated with temperature, salinity, nitrate concentrations, and diversity (viral diversity in the polar and non-viral diversity in the non-polar) (Spearman correlations FDR < 0.01, empty boxes correspond to non-significant correlations). (**C**) Temperature and salinity significantly impact interactome structure in non-polar, while salinity and nitrogen limitation impact most the polar interactome.

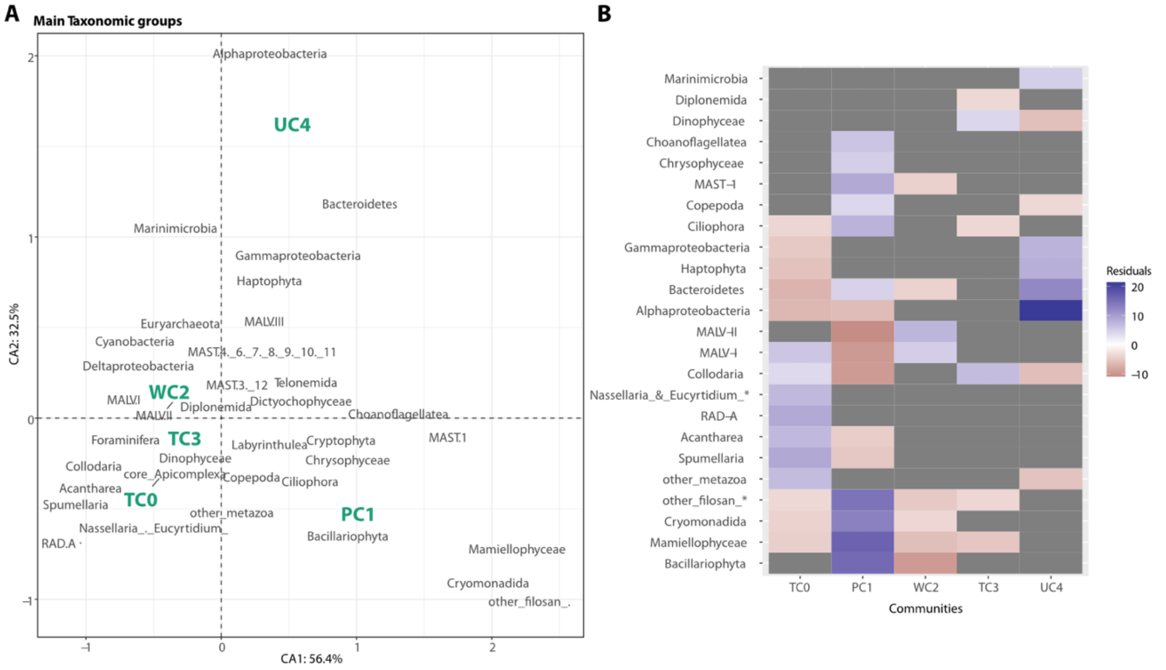


**Fig. S2. The GPI communities differ in their taxonomic composition. (A)** A correspondence analysis reveals how GPI communities (n = 5) relate to each other according to their taxonomic groups in associations. **(B)** Specific lineages are detected as significantly over- (blue) or under- (red) represented within each community via a post hoc analysis for the Pearson's Chi-squared test on the residuals.

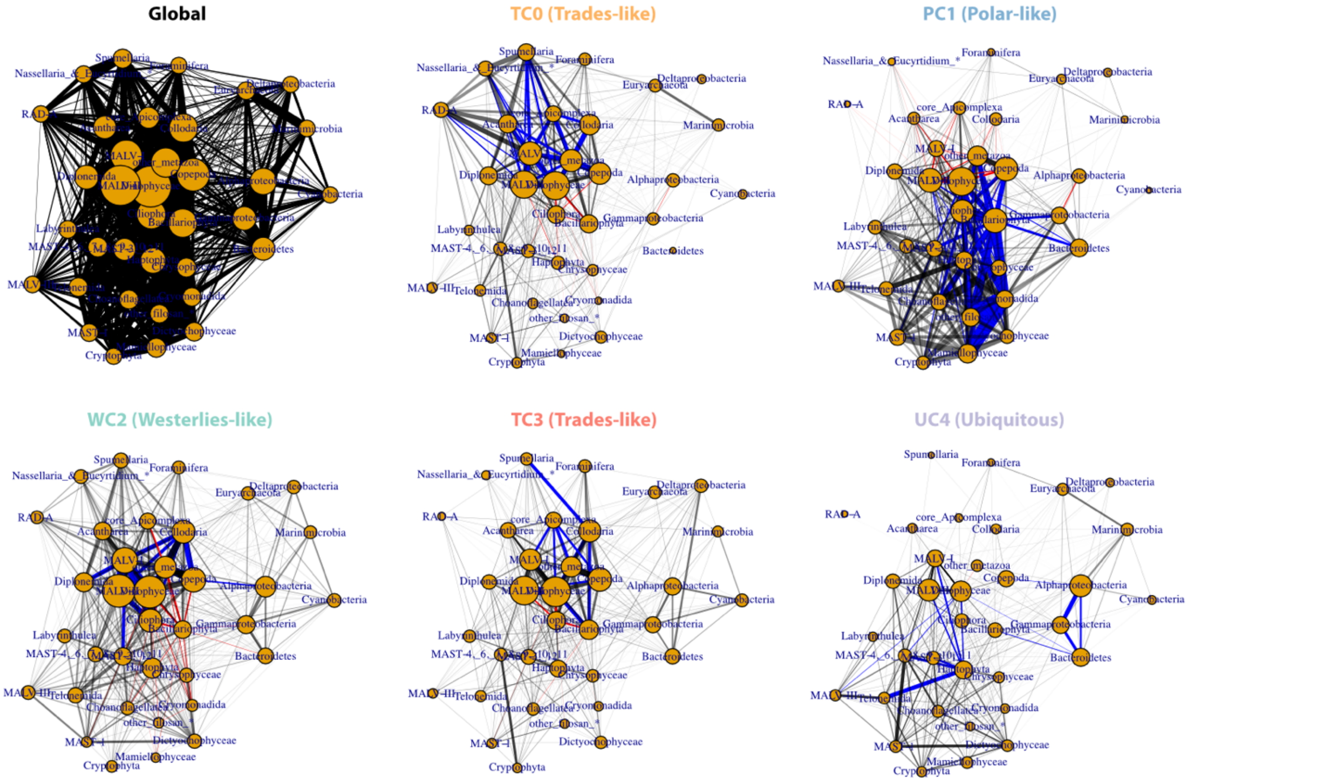


**Fig. S3. The GPI communities display mostly exclusive associations between main planktonic lineages.** A large number of predicted associations are community-specific and several taxa-taxa associations are detected as significantly over- (blue edges) or under- (red edges) represented within each community via a post hoc analysis for the Pearson's Chi-squared test on the residuals.

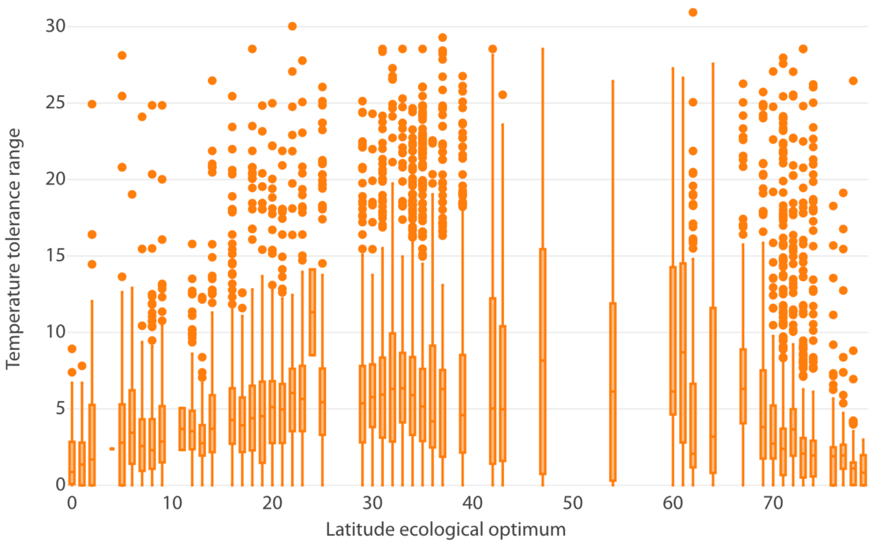


**Fig. S4. Distribution of OTUs temperature tolerance ranges vs. their latitude ecological optima.** The distribution of OTUs temperature tolerance ranges along the latitude ecological optimum is nonrandom, plankton temperature tolerance ranges are smaller towards the pole and the equator.

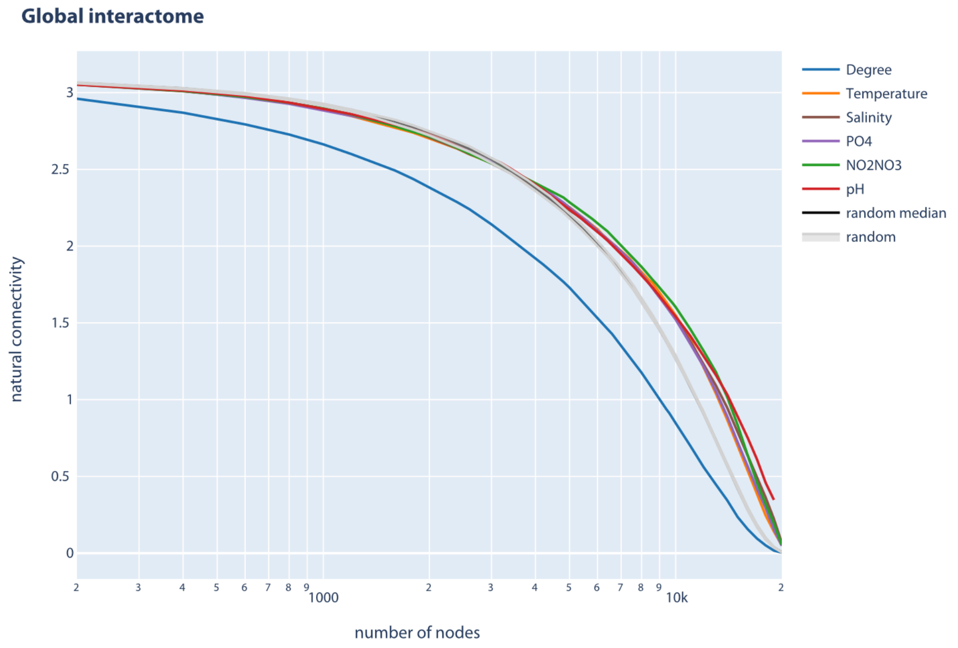


**Fig. S5. Predicted robustness of the GPI at global scale.** Global perturbations of the GPI by distinct environmental tolerance ranges suggest its overall global robustness (estimated by natural connectivity, a graph robustness measure) to environmental changes, as compared to random perturbations (grey curve, n = 100 random perturbations) and to severe perturbations (blue curve, node degree-based perturbations).

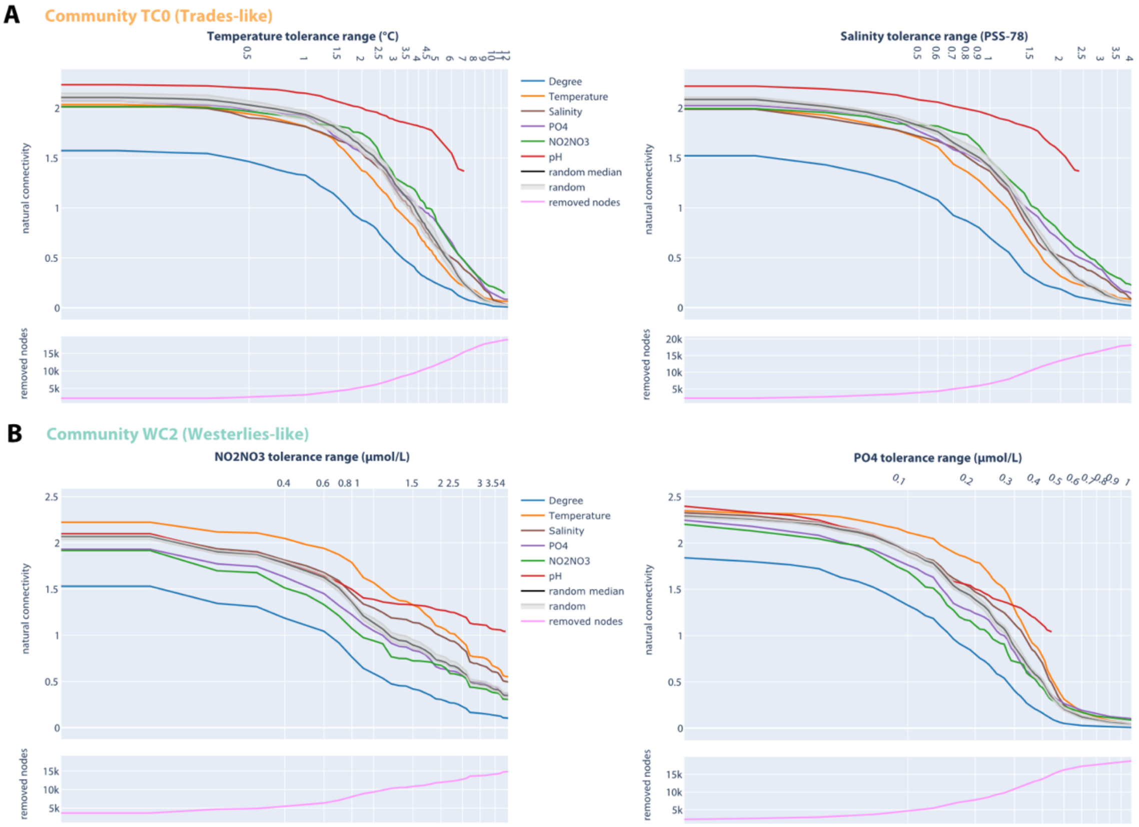


**Fig. S6. Predicted vulnerabilities of the GPI at community scale. (A)** Global perturbations of the GPI by distinct environmental tolerance ranges predict community-specific local vulnerabilities of a Trades-like community interactome (TC0) to temperature (Wilcoxon-rank test, $p = 3.8 \times 10^{-10}$) and salinity ($p = 3.8 \times 10^{-10}$) change. **(B)** Global perturbations of the GPI by distinct environmental tolerance ranges predict community-specific local vulnerabilities of a Westerlies-like community interactome (WC2) to nitrate (Wilcoxon-rank test, $p = 6.7 \times 10^{-9}$) and phosphate change ($p = 5.2 \times 10^{-9}$).

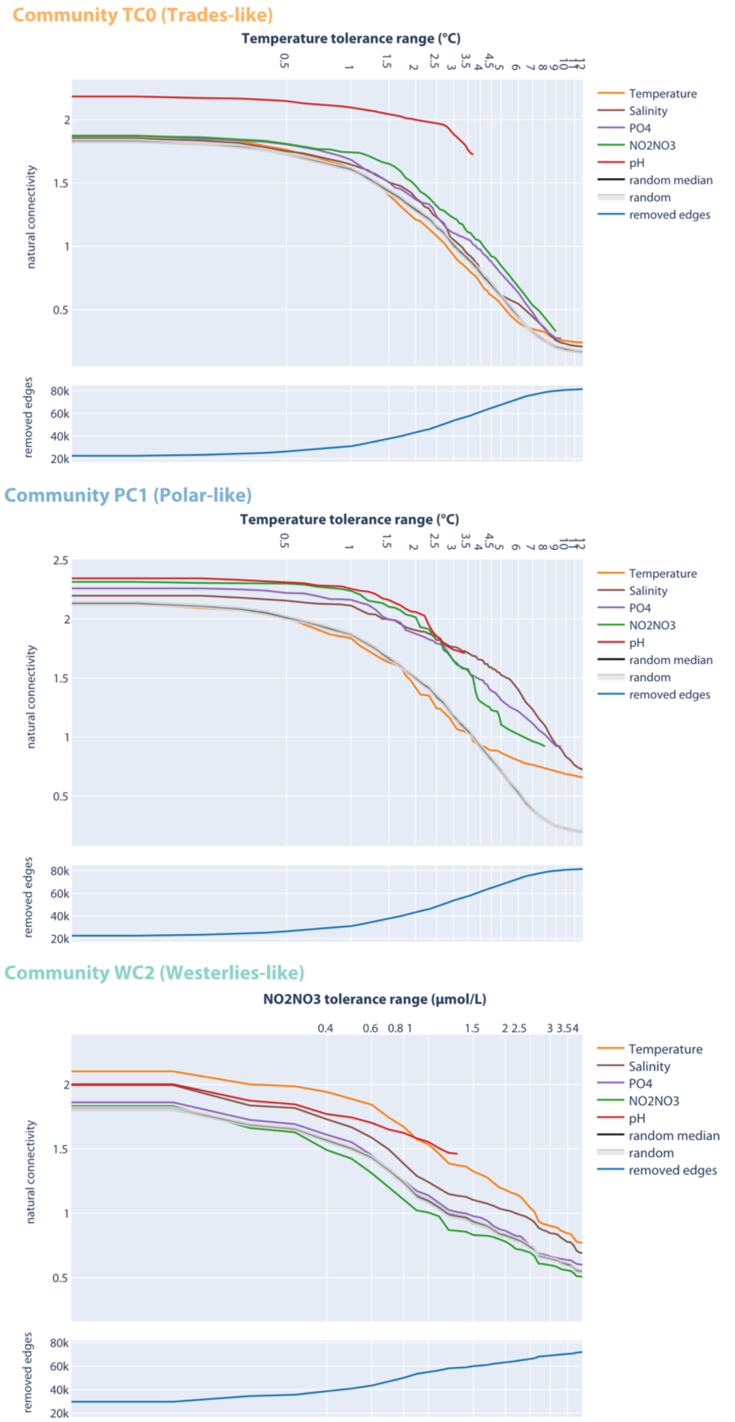


**Fig. S7. Predicted (a)biotic vulnerabilities of the GPI at community scale.** Global perturbations of the GPI by distinct environmental tolerance ranges predict community-specific local vulnerabilities of a Trades-like community interactome (TC0) to temperature (Wilcoxon-rank test, $p = 2.6 \times 10^{-2}$), the Polar-like community interactome (PC1) to temperature ($p = 9.5 \times 10^{-6}$), and the Westerlies-like community interactome (WC2) to nitrate ($p = 6.9 \times 10^{-2}$) change. Here, the GPI is not perturbed by removing nodes but by removing edges instead, from the least to the most overlapping OTU-OTU tolerance ranges, to integrate potential effects of biotic dependencies.

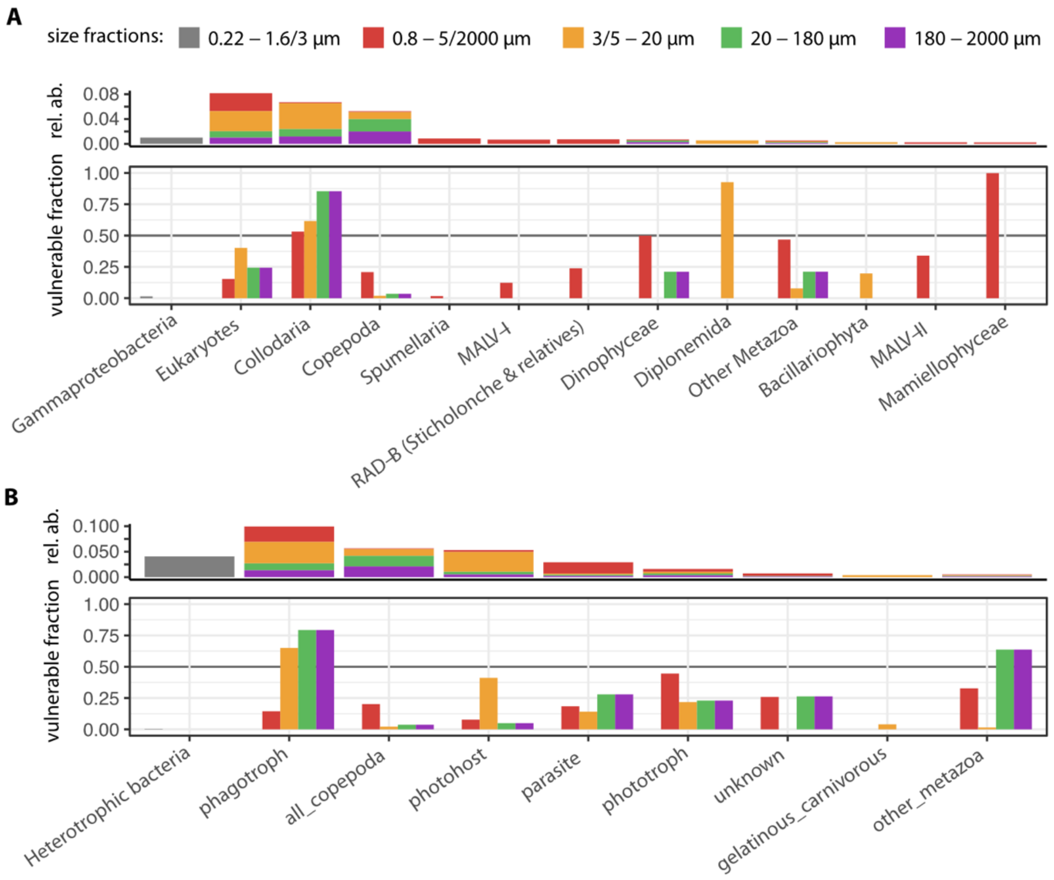


**Fig. S8. Trades-like plankton community lineages predicted most vulnerable to salinity change.** (**A**) Environmental tolerance range perturbations of the GPI predicted Trades-like marine plankton lineages (community TC0) potentially most impacted by temperature variations. **(B)** Grouping these lineages into MPGs predicted associated functions potentially most impacted by salinity variations in the Trades biome. In both panels, the fraction of lineages and MPGs (from 1 for most impacted, to 0 for not impacted) predicted to be impacted by salinity variations are depicted within each size fraction. Plankton lineages (Prokaryotes and Eukaryotes) and MPGs are ordered according to the cumulative mean relative abundance of the corresponding OTUs across size fractions (note that these relative abundances are not directly comparable between size fractions).

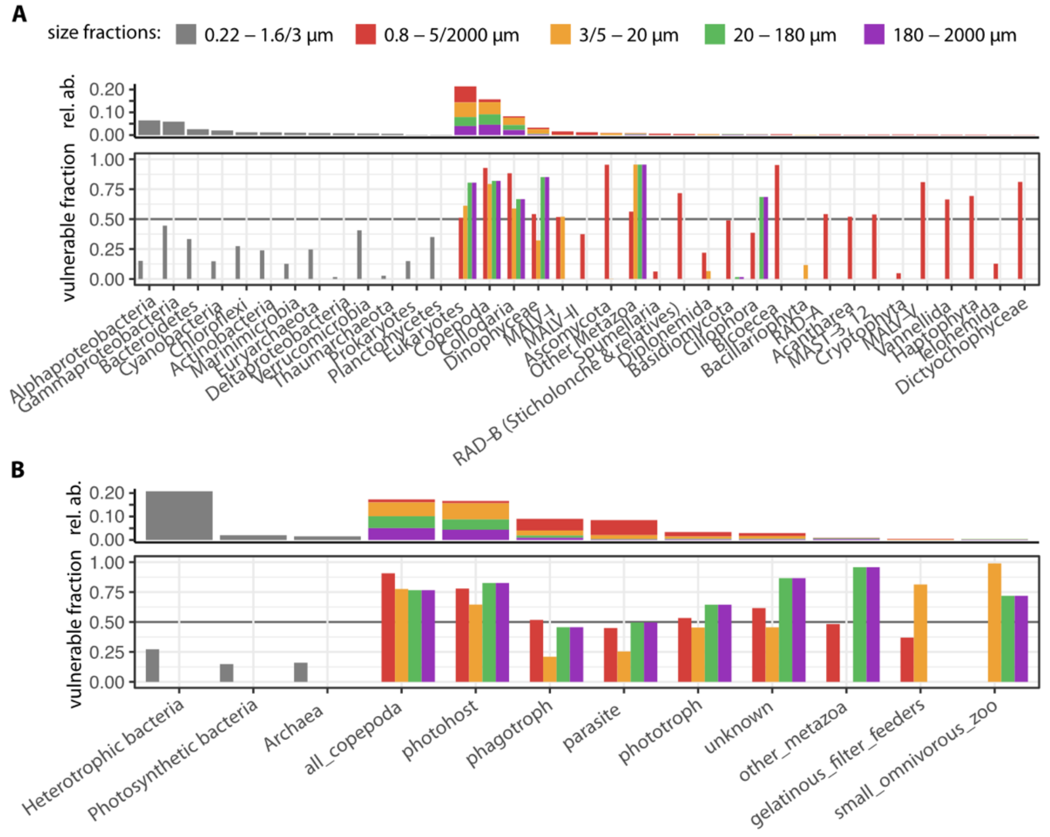


**Fig. S9. Westerlies-like plankton community lineages predicted most vulnerable to nitrate change.** (**A**) Environmental tolerance range perturbations of the GPI predicted Westerlies-like marine plankton lineages (community WC2) potentially most impacted by nitrate variations. **(B)** Grouping these lineages into MPGs predicted associated functions potentially most impacted by nitrate variations in the Westerlies biome. In both panels, the fraction of lineages and MPGs (from 1 for most impacted, to 0 for not impacted) predicted to be impacted by nitrate variations are depicted within each size fraction. Plankton lineages (Prokaryotes and Eukaryotes) and MPGs are ordered according to the cumulative mean relative abundance of the corresponding OTUs across size fractions (note that these relative abundances are not directly comparable between size fractions).

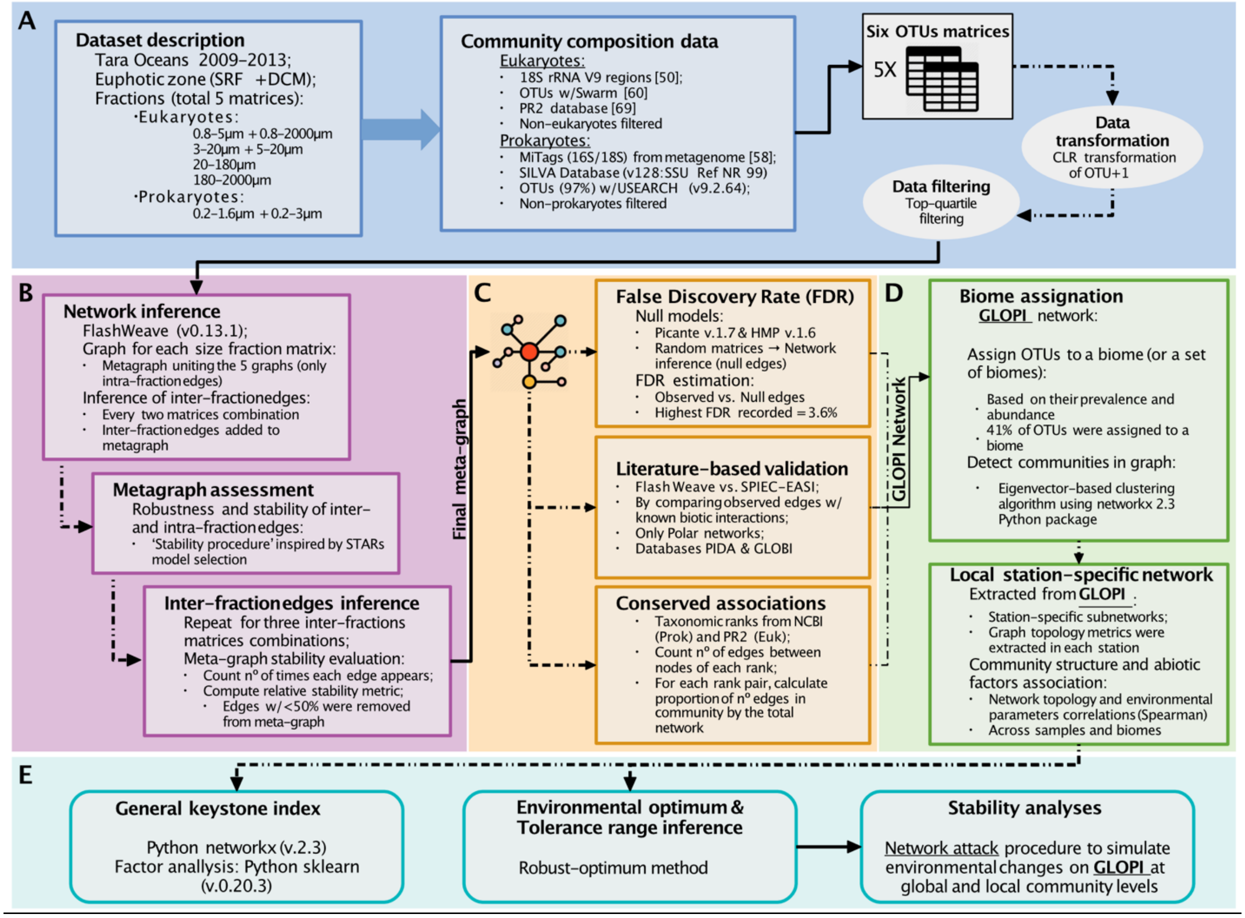


**Fig. S10. The GPI workflow.** (**A**) Data collection, transformation and filtering. (**B**) Networks inference and size fractions integration. (**C**) GPI network false discovery rate estimations via null models and literature validation. (**D**) Biome assignations to GPI OTUs and detected communities, and analyses of station-specific GPI networks. (**E**) GPI general keystone analyses and environmental robustness analyses.

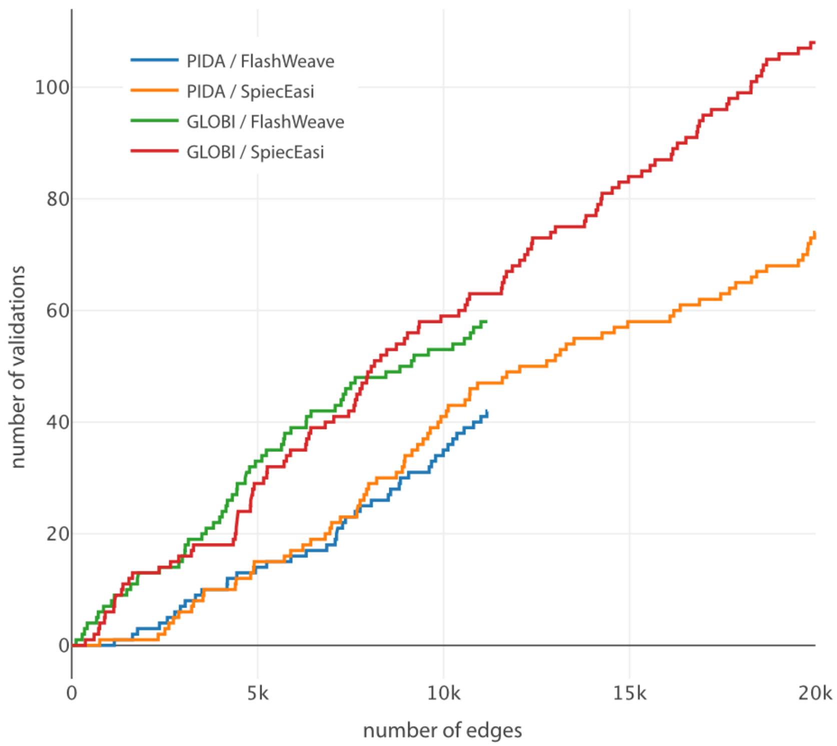


**Fig. S11. Literature-validated fractions of Polar-specific community interactomes.** Networks were reconstructed for Polar samples only (matching size fractions) using both FlashWeave and SPIEC-EASI algorithms. Both methods recovered similar numbers of interactions ranked by their weights at the Genus level (this plot). Similar numbers of interactions were recovered by both methods also at Family and Species taxonomic levels.

### References


1. A. R. Longhurst, in *Ecological Geography of the Sea (Second Edition),* A. R. Longhurst, Ed. (Academic Press, Burlington, 2007), pp. 89-102.
2. A. C. Gregory *et al.*, Marine DNA Viral Macro- and Microdiversity from Pole to Pole. *Cell* **177**, 1109-1123 e1114 (2019).
3. S. Pesant *et al.*, Open science resources for the discovery and analysis of Tara Oceans data. *Sci Data* **2**, 150023 (2015).
4. K. Faust *et al.*, Cross-biome comparison of microbial association networks. *Front Microbiol* **6**, 1200 (2015).
5. F. Mahé, T. Rognes, C. Quince, C. de Vargas, M. Dunthorn, Swarm: robust and fast clustering method for amplicon-based studies. *PeerJ* **2**, e593 (2014).
6. C. Quast *et al.*, The SILVA ribosomal RNA gene database project: improved data processing and web-based tools. *Nucleic acids research* **41**, D590-596 (2013).
7. G. Lima-Mendez *et al.*, Ocean plankton. Determinants of community structure in the global plankton interactome. *Science* **348**, 1262073 (2015).
8. E. Cristóbal, S. V. Ayuso, A. Justel, M. Toro, Robust optima and tolerance ranges of biological indicators: a new method to identify sentinels of global warming. *Ecological Research* **29**, 55-68 (2014).
9. M. F. M. Bjorbaekmo, A. Evenstad, L. L. Rosaeg, A. K. Krabberod, R. Logares, The planktonic protist interactome: where do we stand after a century of research? *ISME J* **14**, 544-559 (2020).
10. J. H. Poelen, J. D. Simons, C. J. Mungall, Global biotic interactions: An open infrastructure to share and analyze species-interaction datasets. *Ecological Informatics* **24**, 148-159 (2014).
11. J. Tackmann, J. F. Matias Rodrigues, C. von Mering, Rapid inference of direct interactions in large-scale ecological networks from heterogeneous microbial sequencing data. *bioRxiv*, 390195 (2018).
12. Z. D. Kurtz *et al.*, Sparse and compositionally robust inference of microbial ecological networks. *PLoS Comput Biol*, 1-25 (2014).
13. S. Mordret *et al.*, The symbiotic life of Symbiodinium in the open ocean within a new species of calcifying ciliate (Tiarina sp.). *ISME J* **10**, 1424-1436 (2016).
14. L. Guidi *et al.*, Plankton networks driving carbon export in the oligotrophic ocean. *Nature* **532**, 465 (2016).
15. X. Yu, M. F. Polz, E. J. Alm, Interactions in self-assembled microbial communities saturate with diversity. *ISME J* **13**, 1602-1617 (2019).
16. M. Striebel, S. Schabhuttl, D. Hodapp, P. Hingsamer, H. Hillebrand, Phytoplankton responses to temperature increases are constrained by abiotic conditions and community composition. *Oecologia* **182**, 815-827 (2016).
17. E. C. Carmack *et al.*, Freshwater and its role in the Arctic Marine System: Sources, disposition, storage, export, and physical and biogeochemical consequences in the Arctic and global oceans. *Journal of Geophysical Research: Biogeosciences* **121**, 675-717 (2016).
18. M. M. Mills *et al.*, Nitrogen Limitation of the Summer Phytoplankton and Heterotrophic Prokaryote Communities in the Chukchi Sea. *Frontiers in Marine Science* **5**, 362 (2018).
19. E. C. Carmack, in *The Freshwater Budget of the Arctic Ocean,* E. L. Lewis, E. P. Jones, P. Lemke, T. D. Prowse, P. Wadhams, Eds. (Springer Netherlands, Dordrecht, 2000), pp. 91-126.

20. D. G. Boyce, B. Petrie, K. T. Frank, B. Worm, W. C. Leggett, Environmental structuring of marine plankton phenology. *Nat Ecol Evol* **1**, 1484-1494 (2017).
21. M. Ardyna *et al.*, Shelf-basin gradients shape ecological phytoplankton niches and community composition in the coastal Arctic Ocean (Beaufort Sea). *Limnology and Oceanography* **62**, 2113-2132 (2017).
22. S. Malviya *et al.*, Insights into global diatom distribution and diversity in the world's ocean. *Proc Natl Acad Sci U S A* **113**, E1516-1525 (2016).
23. R. Logares *et al.*, Disentangling the mechanisms shaping the surface ocean microbiota. *Microbiome* **8**, 55 (2020).
24. D. Righetti, M. Vogt, N. Gruber, A. Psomas, N. E. Zimmermann, Global pattern of phytoplankton diversity driven by temperature and environmental variability. *Sci Adv* **5**, eaau6253 (2019).
25. J. Brate *et al.*, Radiolaria associated with large diversity of marine alveolates. *Protist* **163**, 767-777 (2012).
26. R. Siano *et al.*, Distribution and host diversity of Amoebophryidae parasites across oligotrophic waters of the Mediterranean Sea. *Biogeosciences* **8**, 267-278 (2011).
27. D. K. Stoecker, M. D. Johnson, C. deVargas, F. Not, Acquired phototrophy in aquatic protists. *Aquatic Microbial Ecology* **57**, 279-310 (2009).
28. O. Flegontova *et al.*, Extreme Diversity of Diplonemid Eukaryotes in the Ocean. *Curr Biol* **26**, 3060-3065 (2016).
29. T. Biard *et al.*, Biogeography and diversity of Collodaria (Radiolaria) in the global ocean. *ISME J* **11**, 1331-1344 (2017).
30. T. O. Delmont *et al.*, Nitrogen-fixing populations of Planctomycetes and Proteobacteria are abundant in surface ocean metagenomes. *Nat Microbiol* **3**, 804-813 (2018).
31. T. Biard *et al.*, In situ imaging reveals the biomass of giant protists in the global ocean. *Nature* **532**, 504-507 (2016).
32. R. Derelle, P. López-García, H. Timpano, D. Moreira, A Phylogenomic Framework to Study the Diversity and Evolution of Stramenopiles (=Heterokonts). *Molecular biology and evolution* **33**, 2890-2898 (2016).
33. C. de Vargas *et al.*, Ocean plankton. Eukaryotic plankton diversity in the sunlit ocean. *Science* **348**, 1261605 (2015).
